# Genome-wide mapping of fluoroquinolone-stabilized DNA gyrase cleavage sites displays drug specific effects that correlate with bacterial persistence

**DOI:** 10.1101/2022.10.27.514060

**Authors:** Juechun Tang, Mark P. Brynildsen

## Abstract

Persisters are rare phenotypic variants that are suspected to be culprits of recurrent infections. Fluoroquinolones (FQs) are a class of antibiotic that facilitate DNA damage by stabilizing type II topoisomerases when they are in a complex with cleaved DNA. In *Escherichia coli*, DNA gyrase is the primary FQ target, and previous work has demonstrated that persisters are not spared from FQ-induced DNA damage. Since DNA gyrase cleavage sites (GCSs) largely govern the sites of DNA damage from FQ treatment, we hypothesized that GCS characteristics (*e.g*., number, strength, location) may influence persistence. To test this hypothesis, we measured genome-wide GCS distributions after treatment with a panel of FQs. We found drug-specific effects on the GCS distribution and discovered a strong negative correlation between the cumulative cleavage strength across the chromosome and FQ persister levels. Further experiments and analyses suggested that persistence was not governed by cleavage to individual sites, but rather survival was a function of the cumulative GCS distribution. Together, these findings demonstrate FQ-specific differences in GCS distribution that correlate with persister levels and suggest that FQs that better stabilize DNA gyrase in cleaved complexes with DNA will lead to lower levels of persistence.

## Introduction

Antibiotic treatment failure is a key threat to global public health (1). Beyond the ability of bacteria to acquire resistance toward antibiotics, which refers to growth in the presence of antibiotic concentrations that inhibit susceptible strains, bacteria can exhibit tolerance toward antibiotics, which corresponds to slower death rates in the presence of antibiotics in comparison to susceptible cells (2–4). In addition, hyper-tolerant subpopulations of bacteria called persisters are found in most cultures, where they exhibit death rates that are significantly slower than that of the majority of the population (2). Persisters are phenotypic variants, rather than genetic mutants, that are thought to give rise to recurrent infections, as well as facilitate the development of antibiotic resistance (5–10). For these reasons, it has been postulated that greater understanding of the factors that influence persister survival could reveal counter-measures to improve the treatment of recalcitrant infections (2, 11, 12).

Though multi-drug tolerance can be a feature of some persisters, many persisters exhibit tolerance that is drug specific (7, 13–17). For example, diauxic transitions generated ampicillin (AMP) and ofloxacin (OFL) persisters that were shown to be largely distinct subpopulations (14); amikacin treatment was found to result in far lower persister levels than treatments with AMP or norfloxacin (NOR) (17); and Δ*tisAB* had far fewer ciprofloxacin (CIP) persisters than wild-type, but similar levels of AMP and streptomycin persisters (16). These findings suggest that persistence is often a function of the antibiotic used, which is in contrast to originating from a universal persister formation pathway.

Fluoroquinolones (FQs) are widely used for the treatment of respiratory and urinary tract infections (18), and they are particularly notable in that they can kill both growing and nongrowing bacteria (19); although, their killing capacity wanes in growth-inhibited populations (20, 21). FQs kill bacteria by interfering with the maintenance of DNA topology, targeting primarily DNA gyrase and secondarily topoisomerase IV in *Escherichia coli* (22–24), although it should be noted that topoisomerase IV has been found to be the primary target in some other types of bacteria (*e.g., Streptococcus pneumoniae*) (25). When catalytically active, DNA gyrase produces a staggered double-stranded cut in DNA with 4-bp 5’-overhangs, whose phosphate backbones are covalently linked to Tyr122 of the two GyrA molecules of the enzyme (DNA Gyrase: GyrA_2_GyrB_2_) (26–28). The addition of FQs stabilizes the gyrase-DNA cleaved complexes and inhibits religation, ultimately resulting in DNA breaks that if not repaired lead to cell death (23, 29, 30). In previous work, we demonstrated that persisters to FQs in growth-inhibited populations did not survive due to lack of FQ-induced DNA damage, but rather were able to repair the damage caused by the antibiotic (8, 21, 31, 32). Specifically, FQ persisters were observed to filament and induce the SOS responses, which are canonical indicators of DNA damage (8, 21), and DNA repair mutants had very low FQ persister levels (8). Further work on exponentially-growing cultures demonstrated that FQ persisters in those populations also experienced DNA damage from treatment (33). We later showed that chromosome number (ploidy) was an important variable for FQ persistence, because it defined those individual bacteria that would be proficient with homologous recombination (HR), and thus able to efficiently repair the DNA damage induced by FQs (32). Interestingly, while phenotypic differences in DNA repair have been shown to influence FQ persistence (31, 32), less is known regarding how variables of FQ-induced DNA damage impact persister levels.

Here, we hypothesized that characteristics of FQ-induced DNA damage may influence persistence to FQs. Specifically, we examined whether the number, location, or strength of FQ-stabilized DNA cleavage sites influenced persister survival. We focused on the primary target of FQ in *E. coli*: DNA gyrase (22–24), and developed GCS-seq, which is a chromatin immunoprecipitation (ChIP)-seq method to identify GCSs across the genome that builds upon previous approaches with updates to the sample preparation and sequence analysis pipelines (34, 35). We applied GCS-seq to stationary-phase *E. coli* treated with five different FQs: levofloxacin (LEVO), moxifloxacin (MOXI), norfloxacin (NOR), ciprofloxacin (CIP), and gemifloxacin (GEMI). We identified extensive cleavage throughout the chromosome (20,000+ potential GCSs), and different cleavage patterns between the FQs with respect to the number of distinct GCS, cleavage strength at those sites, and their location. Further, we observed a strong negative correlation between the cumulative cleavage strength for different FQs across the genome and persister levels, and provide evidence that FQ persistence is unlikely to be governed by any individual cleavage sites. These data demonstrate that FQs have both unique and shared GCSs that they stabilize, and suggest that the degree of gyrase cleavage stabilization across the chromosome is a strong predictor of persister levels in growth-inhibited populations.

## Materials and Methods

### Bacterial Strains and Plasmids

All strains and plasmids used in this study are listed in Supplementary Table S1. *E. coli* strain MG1655 was used as wild-type. All other strains used in the experiments, unless indicated, were derived from MG1655. DNA oligonucleotides used for cloning and gene perturbation verification are listed in Supplementary Table S2 (Integrated DNA Technologies, IDT, Coralville, IA, USA).

To chromosomally knock in the strong gyrase cleavage sequence Mu or a scrambled version (MuScr), gBlock gene fragments (IDT, Coralville, IA, USA) containing the desired sequences were first cloned into pKD3 or pKD4 (contain R6Kγ origin) using Gibson Assembly (NEB Gibson Assembly Cloning Kit) and maintained in PIR1 *E. coli* (Invitrogen). The antibiotic resistance cassette and the Mu/MuScr sequence were then amplified from plasmids using PCR and integrated into desired chromosomal loci using the Datsenko and Wanner method (36) with primers indicated in Supplementary Table S2. The same protocol was employed for the construction of C-terminus FLAG-tagged GyrA and its control for GCS-seq experiments. MG1655 Δ*recD* were constructed by P1 transduction (37) from the corresponding mutant in the Keio collection (38). Where indicated, the resistance cassettes were removed using pCP20 following steps as previously described (36, (39). All sequences were confirmed by PCR and sequencing (Genewiz, South Plainfield, NJ).

### Reagents

All media components, chemicals, and antibiotics used in this study were purchased from Sigma Aldrich (St. Louis, MO) or Fisher Scientific (Waltham, MA), unless otherwise noted. In all experiments, autoclaved MilliQ water (18.2 MΩ·cm at 25 °C, Merck Millipore Ltd, Burlington, MA) was used as solvent unless otherwise indicated. Luria-Bertani (LB) medium (Fisher Scientific) was prepared from individual components using 10 g/L tryptone, 5 g/L yeast extract, and 10 g/L NaCl, and LB agar was prepared with 25 g/L BD Difco pre-mixed LB Miller broth and 15 g/L agar. The mixed solutions were then autoclaved at 121°C for 30 min to achieve sterilization. M9 glucose media was prepared using autoclaved 5x M9 minimal salts (33.9 g/L Na_2_HPO_4_, 15 g/L KH_2_PO_4_, 5 g/L NH_4_Cl, 2.5g/L NaCl), 0.1 mM CaCl_2_, 2 mM MgSO_4_, and 10 mM glucose as the sole carbon source. M9 glucose media was filter-sterilized using a 0.22 μm filter (Merck Millipore Ltd, Burlington, MA) upon preparation. For mutant selection and plasmid maintenance, AMP and kanamycin (Kan) were used at a working concentration of 100 μg/mL and 50 μg/mL, respectively, and sterile filtered using 0.22 μm syringe filter (Merck Millipore Ltd, Burlington, MA). Chloramphenicol (Cam) was dissolved in ethanol and used at a working concentration of 25 μg/mL. For mutant selection on plates, the appropriate antibiotic(s) were added into LB agar after it cooled down to about 50°C.

For FQ persistence assays and GCS-seq, stock solutions of 5 mg/mL levofloxacin (LEVO) and 5 mg/mL moxifloxacin (MOXI) in 20 mM NaOH, 1 mg/mL ciprofloxacin (CIP) and 5 mg/mL norfloxacin (NOR) in 0.2 N HCl, and 5 mg/mL gemifloxacin (GEMI) in MilliQ water, were prepared and filter-sterilized using 0.22 μm filters (Merck Millipore Ltd, Burlington, MA). All stock solutions were prepared fresh for each experiment. To determine the antibiotic concentrations used for GCS-seq and persistence assays, the stock solutions indicated above were further diluted to different concentrations (0-10 μg/mL) for FQ survival assays. We selected a working concentration in the concentration-independent region of survival curves where increasing the concentration failed to decrease survival appreciably; except for CIP where the concentration chosen corresponded to the lowest survival observed. A working concentration of 5 μg/mL LEVO, 5 μg/mL MOXI, 1 μg/mL CIP, 5 μg/mL GEMI, and 5 μg/mL NOR were selected. ChIP lysis buffer was prepared using ChIP buffer (sterile 50 mM Tris pH 7.5, autoclaved 150 mM NaCl, sterile 1mM EDTA, 1% Triton X-100) supplemented with 200 U/ml Ready-Lyse lysozyme solution (Lucigen, Middleton, WI) diluted in TE buffer and 1 pill/10 mL Roche cOmplete EDTA-free protease inhibitor cocktail (Millipore sigma, Saint Louis, MO). TBS buffer (50 mM Tris, pH 7.5, 10 mM EDTA) was filter-sterilized using a 0.22 μm filter (Merck Millipore Ltd, Burlington, MA) and kept at 4 °C. Proteinase K solution (20 mg/mL) and RNase (DNase and protease-free, 10 mg/mL) were purchased from Thermo Fisher Scientific (Waltham, MA).

### Fluoroquinolone (FQ) Persistence Assay

Cells were inoculated from glycerol stocks stored at −80 °C into 2 mL LB supplemented with antibiotics for selection as needed and incubated at 37 °C with shaking (250 rpm) for approximately 5 h. Afterward, 20 μL of cultures were added to 2 mL M9 glucose media in test tubes and incubated at 37 °C with shaking for 16 h. Following that overnight growth, cells were inoculated to an OD_600_ ~0.01 in 25 mL M9 glucose media in 250 mL baffled flasks and incubated at 37 °C with shaking for 20 h. After 20 h of incubation, cultures were treated with 5 μg/mL LEVO, 5 μg/mL MOXI, 1 μg/mL CIP, 5 μg/mL GEMI, or 5 μg/mL NOR, respectively, for 5 h. Autoclaved MilliQ water were used as treatment controls. Right before the addition of FQs or water, and at 0.5, 1, 3, 5 h following treatment, 500 μL samples were removed and washed twice. Each wash encompassed centrifugation at 21130 g for 3 min, removal of 450 μL of supernatant, and resuspension of the cell pellet with 450 μL phosphate-buffered saline (PBS). The two wash steps reduced the FQ concentrations by 100-fold, which was below or near the FQ MICs (Supplementary Figure S1). After the washes, cells were serially diluted in PBS, which dropped FQ concentrations further, and plated onto LB agar for CFU enumeration after 16 h incubation at 37 °C.

### Determination of Minimum Inhibitory Concentration (MIC)

MICs of FQs (LEVO, MOXI, CIP, GEMI, and NOR) against strain Mu_origin1_gyrA-FLAG were determined as previously described (39). Briefly, 96-well plates were prepared with varying concentrations of FQs and mixed with a low-density culture (OD_600_ estimated to be ~ 0.0001) that has been diluted in Mueller-Hinton Broth (MHB). OD_600_ values were determined after 16-18 h of incubation in MHB at 37°C. The MIC was defined as the lowest antibiotic concentration at which no cell growth was observed (Supplementary Figure S1).

### GCS-seq: Chromatin Immunoprecipitation using FQs

Strain Mu_origin 1_gyrA-FLAG and FLAG-less controls were used for GCS-seq. Cells were prepared following the same protocol as the FQ persistence assay. Following the addition of FQ, cultures were harvested for ChIP after 30 min of incubation at 37°C with shaking, the beginning of the persister-dominated portion of the survival curve. To harvest cells, cultures were centrifuged at 3220 g for 10 min at 4°C, washed twice with 10 mL ice-cold PBS, and cell pellets were stored at −80 °C for at least 24 h until further processing.

All of the following steps were performed at 4°C. Cell pellets were thawed and resuspended in 2 mL ChIP lysis buffer followed by incubation for ~1 h. After incubation, samples were sonicated for ~10-12 min (10 sec on/10 sec off, 30% amplitude) to yield DNA fragments of ~300-500 bps using Q500 sonicator (500W, 110V with four probe horn assembly, QSONICA, Newtown, CT). Lysates were then centrifuged at 3220 g for 10 min and the cleared supernatants were used for further processing. Twenty μL of supernatants were used as input controls and stored at −20°C in low-bind tubes (Eppendorf, Enfield, CT). For each sample, 50 μL Agarose Anti-FLAG M2 gel pack (Sigma-Aldrich, St. Louis, MO) was prepared by three successive washes with 1 mL TBS buffer and equilibration with 1 mL ChIP Lysis Buffer. Each wash was achieved by centrifugation at 8000 g for 30 sec and removal of supernatant. Nine hundred μL of supernatants were transferred to prepared resins and incubated at 4 °C for 18 h on a tube rotator (Thermo Fisher Scientific, Waltham, MA).

After 18 h of incubation, samples were washed twice with 1 mL ChIP buffer and once in 1 mL TE buffer at room temperature. Each wash was achieved by centrifugation at 8000 g for 30 sec and removal of supernatants. TE buffer was added to both the ChIP-ed samples and thawed input controls to reach a total volume of 200 μL. Five μL of 20 mg/mL proteinase K solution were added to the 200 μL samples, which were then incubated for 3 h at 55°C with occasional vortexing. After 3 h of incubation, samples were treated with 5 μL RNase (10 mg/mL) and further incubated for another 30 min at 37°C. Following that incubation, samples were purified using Qiagen’s PCR Purification kit (Germantown, MD). Note that we added 800 μL PB buffer to each sample and eluted to a final volume of 30 μL in water. Three independent biological replicates were done for each treatment condition. The shearing efficiency and DNA concentration were assessed using a bioanalyzer (Agilent Technologies, Santa Clara, CA) and Quibit 2.0 (Thermo Fisher Scientific, Waltham, MA).

### GCS-seq: SDS-PAGE and Mass Spectrometry Analysis

To validate immunoprecipitation quality, 10 μL of ChIP-ed solutions were removed before the addition of proteinase K, mixed with equal volumes of 2X Laemmli sample buffer (Bio-Rad, Philadelphia, PA), loaded onto SDS-PAGE gels (Mini-PROTEAN Precast Protein Gels, Bio-Rad, Philadelphia, PA), and run at 200V for 40 min following manufacturers’ instructions. Gels were then placed in water for 15 min with shaking at room temperature and stained overnight in staining solution (50% (v/v) methanol, 0.05% (w/v) Coomassie brilliant blue R-250 (Bio-Rad, Philadelphia, PA), 10% (v/v) acetic acid). Subsequently, gels were washed twice in ~ 500 mL boiling MilliQ water for 8 min for distaining. The resulting bands (shown in Figure 1A) were carefully cut and sent for mass spectrometry analyses (Proteomics & Mass Spectrometry Facility, Princeton, NJ). All analyses were performed using an Easy-nLC 1200 UPLC system coupled with an Orbitrap Fusion Lumos (Thermo Scientific, USA). In-gel thiol reduction/alkylation and trypsin Gold (Promega) digestion were performed as in (40). Samples were dried using SpeedVac and resuspended with 21 μL of 0.1% formic acid (pH 3). Two μL of samples were injected per run and loaded directly onto a 45 cm long 75 μm inner diameter nanocapillary column packed with 1.9 μm C18-AQ (Dr. Maisch, Germany). The column temperature was set to be 50 °C and employed one-hour gradient with a flow rate of 300 nL/min (mobile phase compositions: 0.1% formic acid in water and 0.1% formic acid in 80% acetonitrile/water). The mass spectra were collected in data dependent mode. MS scan (positive mode, profile data type, AGC 4e5, Max IT 54 ms, 300-1500 m/z, 120,000 resolution) in the Orbitrap was followed up by HCD fragmentation in the ion trap with 35% collision energy (AGC 1e4, maximum IT of 54 ms, minimum of 5000 ions). Previously fragmented peptides were prevented from repeated fragmentation for 60 sec. Peptides were isolated in a quadrupole (1.2 Da window).

**Figure 1.**
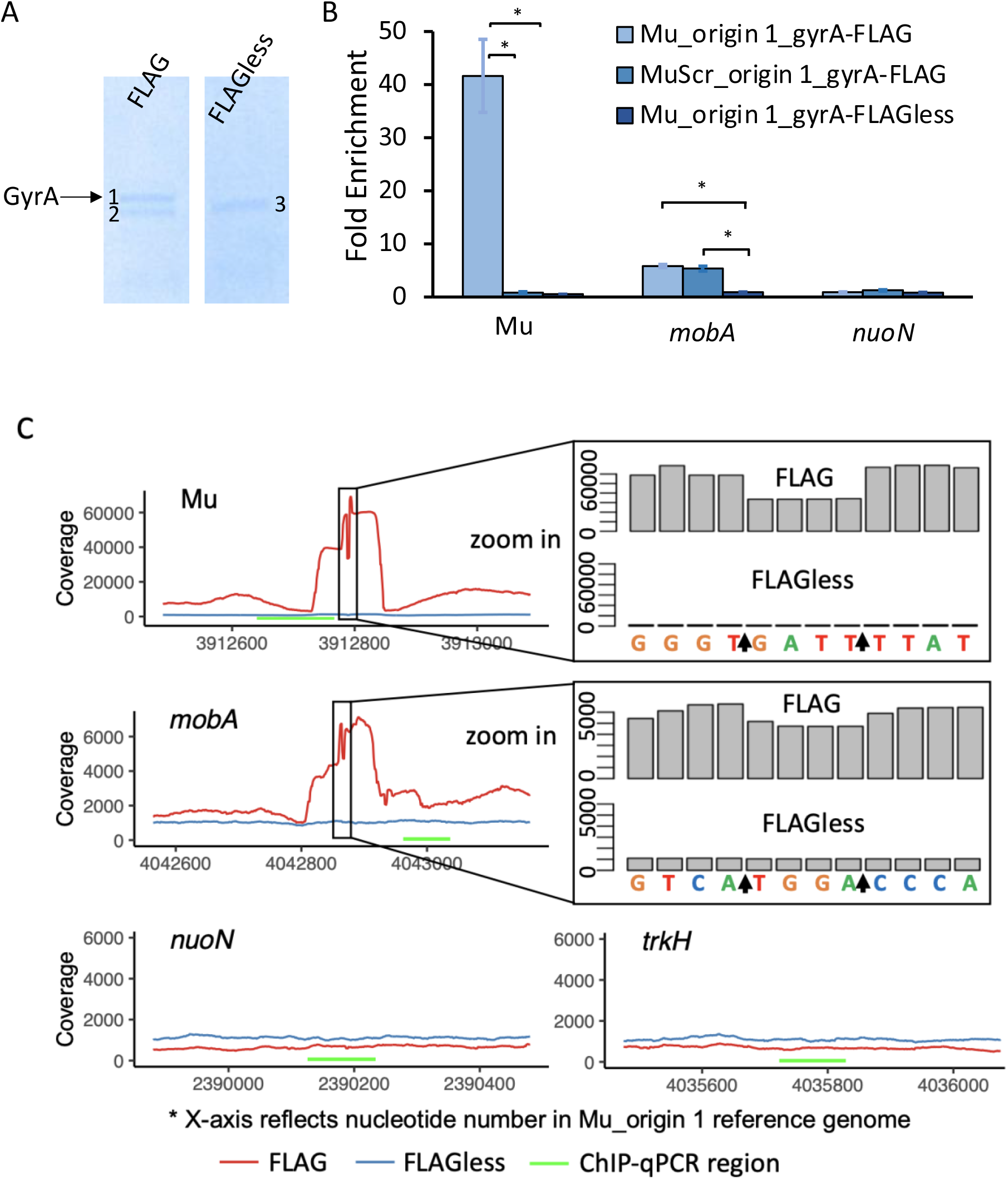
GCS-seq validation in starving cells. FLAG-tagged strains and the FLAG-less controls were immunoprecipitated using anti-FLAG M2 gel following FQ treatment as described in Materials and Methods. (A) Chromatin immunoprecipitated solutions were loaded for SDS-PAGE and three individual bands shown on the gel (indicated as 1,2,3) were excised for mass spectrometry analysis. Left: Strain Mu_origin 1_gyrA-FLAG. Right: Strain Mu_origin1_gyrA-FLAGless. Band 1 showed an overrepresentation of the target protein, GyrA. (B) Fold enrichment of chromatin immunoprecipitated samples at Mu, *mobA*, and *nuoN* sites were measured by qPCR (*n* = 3) and normalized by the *trkH* site. Statistical analysis was performed by comparing between FLAG-tagged strains and the FLAG-less control strains at each site. Data are presented as mean of enrichment ± SEM. Asterisk (*) denotes statistical significance (*p* < 0.05, using two-tailed *t*-tests with unequal variances). (C) Visualization of the coverage depths at Mu, *mobA*, *nuoN*, and *trkH* sites. Local sequences (Reference: Mu-origin 1 strain) are indicated below the coverage track. Arrows indicate cleavage positions. GCSs show a characteristic 4-bp trough. Green bars represent the selected regions where we performed qPCR as shown in (B).

Raw files were searched against UP000000558 *E. coli* database downloaded from UniProt.org. using Sequest HT algorithms (41) within the Proteome Discoverer 2.4 suite (Thermo Scientific, USA) using 10 ppm MS1 and 0.4 Da MS2 mass tolerances. Carbamidomethylating of cysteine was set as a fixed modification, and oxidation of methionine and acetylation of protein N-termini as dynamic modifications. A maximum of 2 missed trypsin cleavages were allowed. Scaffold (version Scaffold_5.1, Proteome Software Inc., Portland, OR) was used for protein identifications.

### GCS-seq: Quantitative PCR (qPCR) Validation

ChIP-ed samples and their respective input controls were assessed by qPCR at four selected loci using primers listed in Supplementary Table S2. The primer sets and exponential amplification range were assessed to ensure amplification efficiencies around 100% (+/− 10%). For each reaction, 0.5 μL of 10 μM forward and reverse primers were mixed with 10 μL of 2X Power SYBR-green master mix (Applied Biosystems, Waltham, MA) and diluted to a total volume of 19 μL with RNase-free H_2_O. One μL of ChIP-ed DNA or input controls (diluted 100-fold to reach concentrations between ~0.1 −5 ng/μL) were added to each well and mixed in the MicroAMP fast optical 96-well reaction plate (Thermo Fisher Scientific, Waltham, MA). The loaded plate was amplified for 40 cycles via the ViiA 7 real-time PCR system (Thermo Fisher Scientific, Waltham). For each biological replicates (n = 3), three technical triplicates were performed. The fold enrichment was calculated as:

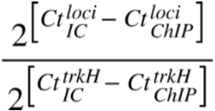

where *Ct*, subscript *IC*, and subscript *ChIP*, represent cycle threshold values, input controls, and ChIP-ed DNA, respectively, whereas *trkH* was used as an unenriched control site for normalization.

### GCS-seq: Library Preparation

Following immunoprecipitation, we used ChIP-ed samples and their respective input controls for library preparation using Accel-NGS 1S plus DNA library kit (Integrated DNA Technologies, formerly Swift Biosciences, Ann Arbor, MI) following manufacturers’ instructions. We used ~20 ng per sample for library preparation. For the clean-up steps, we used SPRIselect purchased from Beckman Coulter (Indianapolis, IN). Swift Unique Dual Indexing Primers (X9096, Swift Biosciences, Ann Arbor, MI) were used for PCR indexing. The samples were mixed at equal amounts (~50 ng) to generate pooled libraries and stored at −20°C. The kit allows library construction from both single and double stranded DNA, which circumvents potential sequencing issues originated from the poor ligation of dsDNA adaptors due to the covalently linked 5’end of DNA and Tyr122 on GyrA (42).

### GCS-seq: DNA Sequencing and Mapping

Sequencing was performed on an Illumina NovaSeq 6000 platform (flow cell SP 100nt Lane, 2 x 50 bp) at Princeton Genomics Core Facility (Princeton, NJ). Reads qualities were assessed with FastQC (43). Demultiplexed data was uploaded to galaxy.princeton.edu. Within Galaxy, sequencing reads were trimmed using Trim Galore! (Galaxy Version 0.6.3) and mapped to Mu_origin 1 genome using BWA-MEM (version 0.7.17.1) (44). Mu_origin1 genome (GEO: GSE206610) was constructed by the insertion of the strong gyrase cleavage sequence Mu into the intergenic region between *pstS* and *glmS* (3911700-3912940) using *E. coli* MG1655 genome (NCBI NC_000913.3) as the background strain. The resulting BAM files were filtered to retain reads that had mapping qualities >=20. IGV (45) was used for visualization.

### GCS-seq: Identification of GCSs

Read depth at each position was calculated using SAMtools depth (46) taking the filtered BAM files as input. The maximum number of considered alignments was set to 100000 to prevent depth saturation with the default upper boundary (default = 8000). For GCS calling, we developed an algorithm that searches for the 4-bp gap pattern using the sequencing coverage depth as the sole input.

The algorithm depends upon a single ‘cut-off’ hyperparameter ***α*** that is shared for all drug treatments, which essentially serves as a threshold of GCS strength. Specifically, for each drug treatment, the input is the coverage at each nucleotide position for the ChIP-ed sample and its respective input control (two arrays of length equivalent to the genome length, ~4.6M). We denote the normalized coverage (normalized by total coverage) as *norm* and *IC_norm*. The algorithm determines whether position *i* is identified as a gyrase cleavage site by checking if *norm/IC_norm* satisfies the following two local conditions for all three replicates: (for convenience, we denote *norm/IC_norm* by function *f*)

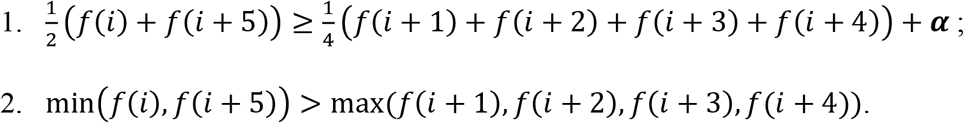

We reason that both conditions (1 and 2) are necessary for identification of gyrase cleavage sites based on the mechanism of action of DNA gyrase. Specifically, using ssDNA library preparation and FQ to immobilize DNA gyrase on DNA, 4 bp 5’ overhangs should be covalently attached to GyrA, and thus those single strands will not be sequenced, which will produce 4 bp troughs in the coverage depth (34). We note that *norm/IC_norm* can occasionally trigger division by 0 errors. In implementation, a small value of ***ε*** = 1*e* − 11 was added to the denominators and parametric analysis suggested that the called cleavage sites were not affected by the choice of *ε* (1e-11 to 1, Supplementary Figure S2).

To select ***α*** and quantify algorithm performance, we ran the algorithm on both FLAG-tagged samples and FLAG-less samples (as a ground-truth dataset that does not exhibit gyrase cleavage) as a function of ***α***.We selected the minimum value of ***α*** that satisfies: (for convenience, we denote the number of GCSs as *n*)

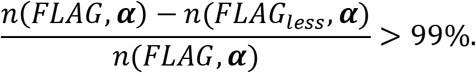

We further define the cleavage strength of j^th^ independent measure at location *i* as:

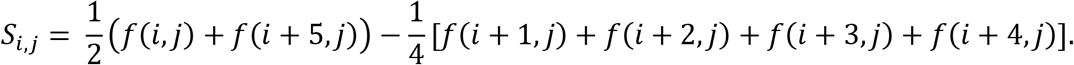

Thus, the average cleavage strength at location *i* is: 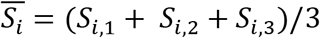.

We note that the expected number of sites identified over a 4.6 Mbp genome with an ***α*** = 0 would be ~1300, if cleavage strengths at each position are independent and identically distributed random variables 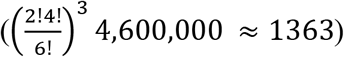

### Motif Construction

For motif construction and visualization, 18 nt long sequences surrounding GCSs (from position *i* − 6 to *i* + 11) were extracted and used for the construction of a position-specific score matrix (PSSM). For a weighted position-specific score matrix the average cleavage strengths of the GCSs were taken into consideration. Specifically, a GCS with relative cleavage strength 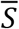 was counted *S*′ times (*S*′ is rounded down from 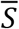 to the nearest whole number) in the weighted position-specific score matrix. PSSM was constructed using Bio.motifs from Biopython package (47). Logos were visualized by WebLogo (48).

### Mu Scrambled Sequence Generation

To construct an uncleavable MuScr control sequence (Supplementary Table S2), 130 nucleotides surrounding the GCSs (from i-63 to i+66) were used for motif construction following steps in the section above. We used the GCSs generated in nutrient-replete conditions as an initial guide (34) and then confirmed the score with data generated in this study. The core of the Mu sequence containing the strong GCS (69-nucleotide long) was scrambled multiple times and scanned using the constructed PSSM. The sequence that showed the lowest log-odds score was selected as a non-cleavable candidate and confirmed by qPCR.

### Generation of Genome Atlas

The genome atlas plot was generated by DNAPlotter (Artemis companion program) (49). The track for each FQ treatment was prepared by custom Python script before imported into DNAPlotter.

### Generation of GCSs Distribution Plot

Mu_origin 1 genome was evenly divided into 10 non-overlapping bins (A to J). All bins contained 464289 nucleotides except bin 10, which contained 3 more nucleotides. The number of GCSs and the cumulative cleavage strength within the range of each bin were represented as bar plots.

To determine if GCS distribution exhibits macrodomain bias, the chromosome was binned based on macrodomain boundaries (Supplementary Table S3) (50). Due to varying length of each macrodomain, the number of GCSs and cleavage strength within each macrodomain were normalized with respect to expected number of GCSs and cleavage strength for visualization purposes (based on a null hypothesis of even distribution of GCSs and cleavage strength across the chromosome).

### RNA-seq: Total RNA Extraction

Strain Mu_origin 1_gyrA-FLAG was used for RNA-seq. The cells were prepared and cultured to stationary-phase following the same procedure as GCS-seq. After 20.5 h of incubation in M9 glucose media, 50 μL of culture was mixed with 100 μL of RNAprotect bacteria reagent (Qiagen, Germantown, MD). Cells were lysed using 1 mg/mL lysozyme dissolved in TE buffer and purified using RNeasy Mini Kit with on-column DNA digestion with DNase I (Qiagen, Germantown, MD) following manufacturers’ instructions. The RNA quality was assessed with a bioanalyzer (Agilent Technologies, Inc, Santa Clara, CA) and Quibit 2.0 (Thermo Fisher Scientific, Waltham, MA).

### RNA-seq: Library Preparation and Sequencing

RNA library preparation was performed by Princeton Genomics Core Facility (Princeton, NJ). Briefly, rRNA depletion was performed using RiboCop rRNA Depletion Kit (Lexogen, Greenland, NH). The Apollo 324 NGS Library Prep System was used with the PrepX RNA-seq library preparation kit (Takara Bio, San Jose, CA). Sequencing was performed on Illumina NovaSeq 6000 platform (SP 100nt Lane, read length 2 x 50 bp) and demultiplexed data was uploaded to Galaxy (galaxy.princeton.edu) for downstream processing.

### RNA-seq: Associate GCSs with Gene Expression Level

Within Galaxy, the reads qualities were assessed and mapped to Mu_origin 1 as reference genome using STAR (51). Reads mapped to each gene were quantified using featureCounts (Galaxy version 1.6.4+galaxy1) (52). The following analysis was performed in R. To examine if gene expression level correlated with gyrase cleavage activity, reads were first normalized by the gene length and further normalized by the sum of the normalized reads. Following gene expression normalization, high expression and low expression genes (normalized expression mean >400 or normalized expression mean <1.5 on average, respectively) were identified. Overall, 404 genes were categorized as low expression genes and 382 genes were identified as high expression genes, which reflects approximately bottom and top 10% of the gene expression distribution. Number of distinct GCSs or the cumulative strength of cleavage in the upstream, downstream, 5’-end, and 3’-end regions of all genes, highly-expressed genes, and genes with low expression were determined, where upstream, downstream, 5’-end, and 3’-end regions were defined as within 300 bp upstream of the start codon, within 300 bp downstream of the stop codon, within 300 bp downstream of the start codon, and within 300 bp upstream of the stop codon, respectively.

### PCA Analysis

Data matrix of cleavage strengths corresponding to the union of GCSs was used as input to PCA analysis. A pseudo cleavage strength using the same formula as above was calculated for sites that were not identified as GCSs, which was bounded by a value of 0. Outlier values (5 out of 23277 GCSs, due to division by 0) were removed. This resulted in a 23272 × 15 matrix M since there was a total of 23272 GCSs identified and 15 samples (5 FQs × 3 biological replicates). PCA was performed after standardization and scaling by unit variance.

### Clustering of FQ-stabilized GCSs

The union of all identified GCSs (same as the input for PCA analysis) or the shared GCSs across different treatment conditions were hierarchically clustered based on their cleavage strengths using R package ComplexHeatmap (53). The heatmap with dendrogram was generated using quantile break color schemes such that each color represents an equal proportion of the data. Complete linkage was used for hierarchical clustering.

### Statistical Analysis

At least three independent biological replicates (*n*) were performed for each experiment. Statistical details of each experiment can be found in the corresponding figure legends. Data are presented as mean ± SEM where suitable. For qPCR enrichment analysis (Figure 1B) and persistence assay (Figure 5, Figure 6, Supplementary figure S11 and S12), significance was assessed using two-tailed student’s t tests with unequal variance for pairwise comparisons. For cleavage strength distribution across different MDs (Figure 3B), normalized values were compared with a value of 1.0 (one-tailed t-test). To assess the cleavage strength with respect to gene expression level (Figure 3C) and cleavage strength across the genome (10 bins, Supplementary Figure S9A), statistical analyses were carried out using one-way analysis of variance (ANOVA) followed by Tukey’s post hoc test. For plots depicting number of GCS distributed across MDs (Supplementary figure S8), across 10 bins (Supplementary figure S9B), and regions related to genes of varying expression level (Supplementary figure S10), exact binomial tests were performed to determine the effects of location (MD, location in the chromosome, and proximity to genes of expression level) on the number of distinct GCSs. For the above cases, an adjusted p value ≤ 0.05 was considered statistically significant.

## RESULTS

### ChIP-qPCR validation of sites with and without GCSs in non-growing *E. coli*

We sought to quantify FQ-stabilized GCSs at the genome-scale in stationary-phase *E. coli*, because such cultures have high persister levels and previous studies had established that FQ persisters in those cultures experienced DNA damage from treatment (8, 21, 31). To do this, we developed GCS-seq, a ChIP-seq approach based on two previous studies that have been used for mapping of topoisomerase binding and cleavage sites in the *E. coli* genome in nutritive environments (34, 35). Specifically, GCS-seq adapted two key features from NorflIP (35) and Topo-Seq (34). First, we used FQs as fixation agents, just as NOR was used in NorflIP (35), and CIP or oxolinic acid (a precursor to the fluoroquinolone class (54)) were used in Topo-Seq (34). Second, we applied single-strand DNA adaptors during library preparation that allowed only DNA fragments with free 3’-ends to be sequenced as was done in Topo-Seq (34). These two key experimental techniques allowed for immunoprecipitation of only gyrase-bound DNA fragments and a characteristic 4-bp trough in read counts at GCSs.

For immunoprecipitation, we chromosomally tagged *gyrA* with a FLAG-tag as in Topo-Seq and NorflIP (34, 35). As controls, we constructed FLAG-less strains that were used in our analysis pipeline to estimate the false-positive rates (Materials and Methods), whereas Topo-Seq used untreated samples (no drug treatment). In addition, the Topo-Seq algorithm uses the 3’-end counts as input and identifies sites where the *i* and *i* + 5 positions both show higher counts in gyrase-poisoned samples compared to the untreated samples (both normalized by input controls) (34). Here, the GCS-seq algorithm uses the sequencing coverage depth as input and searches for 4-bp troughs (Materials and Methods), which is a biological constraint given the catalytic mechanism of DNA gyrase and the sequencing steps used.

To assess whether GCS-seq could be successfully applied to starved *E. coli*, we sought to perform ChIP-qPCR on sites that were previously determined to contain strong or undetectable cleavage, as positive and negative controls, respectively. To accomplish that, we inserted a 1241 bp sequence derived from bacteriophage Mu, which was previously found to experience strong DNA gyrase cleavage (34, 55, 56), close to the origin of replication in wild-type *E. coli* MG1655. We also designed a scrambled equivalent of the Mu site (MuScr), which was expected to eliminate cleavage. Further, we considered *mobA*, which previous work identified to contain a GCS, as well as *nuoN* and *trkH*, which had not been found to contain GCSs (34). For the initial gyrase fixation agent, we used LEVO at a concentration of 5 μg/mL (~80-fold the minimum inhibitory concentration (MIC)) (Supplemental Figure S1), which was chosen based on the concentrations used in persistence assays (32).

Following immunoprecipitation, SDS-PAGE consistently showed 2 separate protein bands of ~100 kDa in the FLAG-tagged strain and 1 band in the FLAG-less control. We note that the lower band from the FLAG-tagged strain ran to the same position as the band from the FLAG-less strain (Figure 1A). To assess the biological identity of the immunoprecipitated proteins, we excised the bands and sent them for mass spectrometry analysis. Mass spectrometry confirmed that GyrA was the most enriched protein in the top-band, whereas the lower-band was mostly comprised of AceE, which is a component of pyruvate dehydrogenase (57, (58) and likely represents nonspecific signal from immunoprecipitation. Quantitative PCR of immunoprecipitated samples using *trkH* as a normalization standard demonstrated that the Mu site was indeed a strong GCS with >40-fold enrichment, whereas such enrichment was completely abolished with MuScr (Figure 1B). Further, *mobA* showed significant enrichment in both FLAG-tagged strains, either with Mu or MuScr integrated, but not the FLAG-less control, whereas enrichment was not observed for *nuoN* in any strain (Figure 1B). These data demonstrated that the ChIP steps of GCS-seq enabled GCSs in stationary-phase bacteria to be identified with qPCR.

### Sequencing shows characteristic 4-bp gap at GCSs

After qPCR validation, we applied the sequencing step (Materials and Methods) of GCS-seq to the immunoprecipitated DNA. Corresponding to the qPCR results, we observed characteristic 4-bp gaps at Mu and *mobA* cleavage sites in FLAG-tagged strains. That pattern was not observed at corresponding sites in the FLAG-less control strain or at control sites (*nuoN* and *trkH*) in the FLAG strain (Figure 1C). These data suggested that we were able to identify sequencing sites that are known to contain and not contain GCSs with the modified procedures described, and that the 4-bp sequencing troughs were readily observable.

### GCS-seq identifies GCSs in starving *E. coli* with high precision

Due to the biological significance of the 4-bp trough, we elected to develop an algorithm that used that biological insight to identify GCSs from the FQ-treated ChIP-seq data (Materials and Methods). Briefly, the algorithm scans the genome comparing counts (normalized with input control, both scaled by their respective total coverage) from nucleotide *i+1* to *i+4* (4 nucleotides of the trough) to those of *i* and *i+5* (2 boundary nucleotides). A GCS is identified at position *i* if, (1) the counts at *i* and *i+5* exceed all of the counts from position *i+1* to *i+4*, (2) the average of the counts of the boundary nucleotides exceeds the average of the counts of the trough nucleotides by a threshold, defined by the parameter ***α***, and (3) conditions 1 and 2 are satisfied in all replicates. The parameter ***α*** can be greater than or equal to 0, and we sought to examine how its value impacted the number of GCSs identified in the FLAG-tagged samples compared to the FLAG-less controls. Specifically, we expected to observe a higher number of GCSs in the FLAG-tagged samples and a close-to-baseline number of sites in the FLAG-less strain. Indeed, we observed far fewer GCSs in FLAG-less strain compared to FLAG-tagged samples for all of the five FQs tested. For example, for LEVO treated sample and ***α*** = 0, we observed over 20,000 GCSs in FLAG-tagged samples, whereas approximately 700 were identified in the FLAG-less control (Supplementary Figure S3AB), which was of the same order of magnitude that would be expected at random (Materials and Methods). As ***α*** was increased, the number of GCSs identified in the FLAG-less control fell precipitously, whereas the decline was far slower in the FLAG-tagged sample (Supplementary Figure S3AB). If it is assumed that at each ***α***, the number of GCSs identified in the FLAG-less control reflects the number of GCSs erroneously identified in the FLAG-tagged sample (false positives), at ***α*** ≥ 0.04, this algorithm achieved precisions over 99% for all five FQs (Supplementary Figure S3C). Manual inspection of GCSs selected at random from those identified at those precision levels in FLAG-tagged samples compared to the same nucleotides in the FLAG-less controls confirmed that they had the expected trough structure of GCSs. Further, Mu and *mobA* sites were identified as GCSs, which was consistent with qPCR and previous work. Altogether, we leveraged biological knowledge of the FQ mechanism of action to develop an algorithm to identify GCSs from ChIP-seq data of FQ-immobilized DNA gyrase-DNA cleaved complexes that exhibits high precision and accurate identification of qPCR-verified GCSs.

### GCS distributions vary based on the FQ used

We applied GCS-seq to quantify GCS distributions of stationary-phase *E. coli* treated with five distinct FQs; MOXI, LEVO, GEMI, CIP, and NOR, at concentrations that would be used for persistence assays (Supplementary Figure S4 and S5). Notably, each FQ was used at a concentration that was ≥ 20-fold its MIC (Supplementary Figure S1). For all FQs, significantly fewer GCSs were identified in the FLAG-less samples, which declined to near 0 as ***α*** was increased. We observed that increasing ***α*** beyond 0.04 only had marginal effects on precision, while the number of GCSs continued to decline due to the more stringent threshold (Supplementary Figure S3). Therefore, we selected ***α*** = 0.04 as the universal threshold for all five FQ treatments; although, it should be noted that the algorithm yielded a consistent order in the number of distinct GCSs (MOXI>LEVO>GEMI>CIP>NOR) over a wide range of ***α*** (0.03 to 0.20, Supplementary Figure S6). In total, GCS-seq identified 14504, 12283, 11644, 6634, and 1551 GCSs in MOXI-, LEVO-, GEMI-, CIP-, and NOR-treated cultures, respectively (Figure 2A). Among all the 23277 identified GCSs (union of all five FQ-treated GCSs), 956 were shared among all five FQs, and each FQ contained GCSs not observed with other FQs (Figure 2B). For a more quantitative comparison of the similarities and differences between the GCS distributions, we performed principal component analysis (PCA) on the cleavage strengths of GCSs identified (Materials and Methods). We found that ~75% of the variation in the data could be explained by the first three principal components and that the different FQ treatments were readily separable in principal components space (Supplementary Figure S7A-D). To complement PCA, we performed unsupervised hierarchical clustering, which revealed clustering based on the FQ used and showed an overall trend where MOXI-treated cultures tended to experience stronger cleavage and NOR-treated cultures tended to show weaker cleavage among FQs used (Figure 2C). We note that the heatmap captured the cleavage strength across the union of all potential GCSs, and we also considered whether clustering of the shared GCSs would reveal anything further. However, clustering based on the cleavage strengths of the 956 shared GCSs produced similar results (Supplementary Figure S7E), where MOXI tended to exhibit stronger cleavage and NOR weaker cleavage among the FQs considered. Interestingly, the heatmaps also revealed deviations from this general pattern, where MOXI did not uniformly produce the greatest degree of cleavage, and NOR did not uniformly produce the weakest degree of cleavage, but rather, site and strength differed to varying levels between the different FQs (Figure 2C and Supplementary Figure S7E). However, it is worth noting that the degenerate consensus motif identified previously for exponential-phase cultures, where G and C are highly abundant at positions *i* and *i+4* (34, 59, 60), was also found to be true for the stationary-phase cultures and all FQs considered here (Figure 2D).

**Figure 2.**
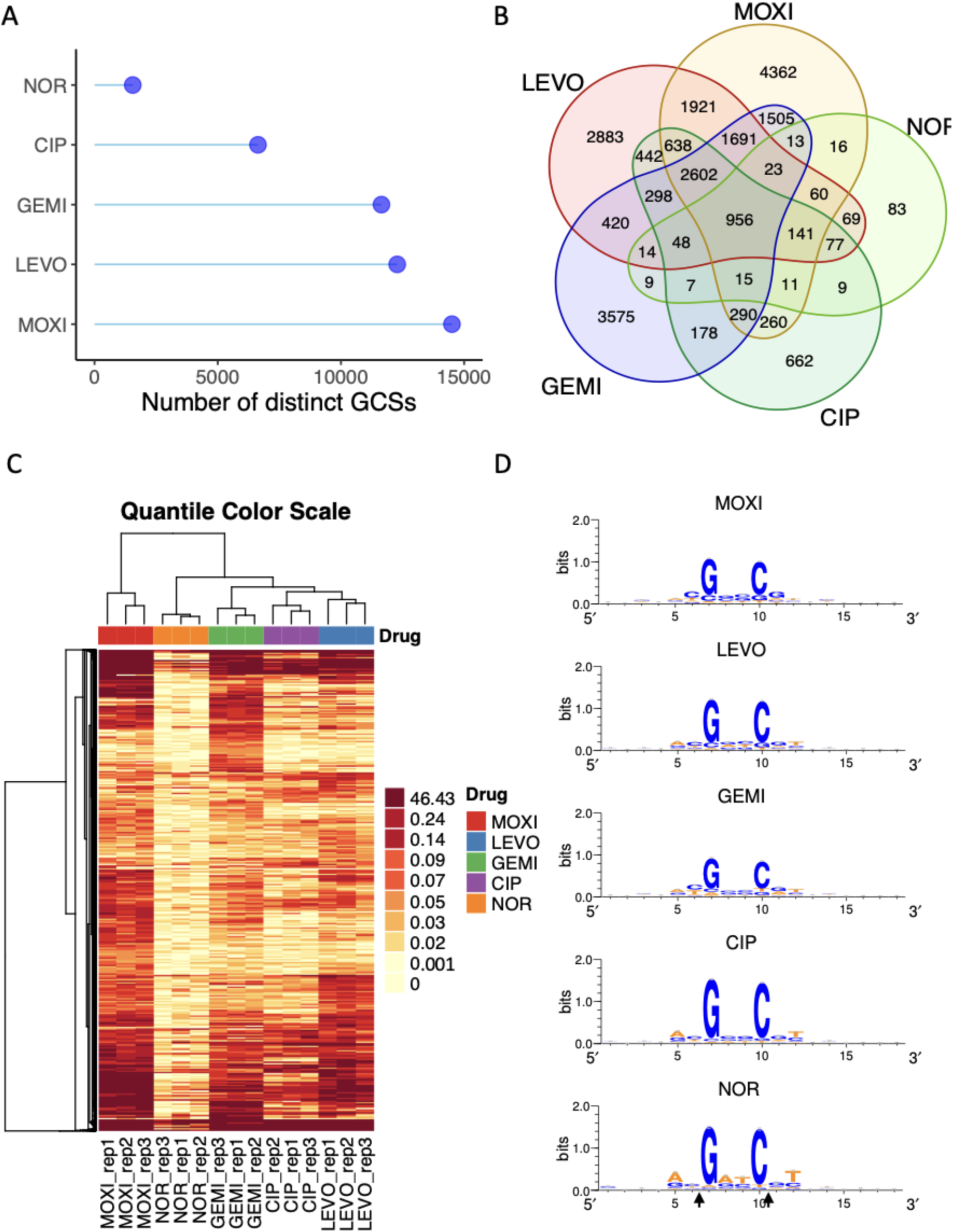
GCS-seq identified distinctive GCS distribution for FQs. (A) Distinct number of GCSs found for MOXI, LEVO, GEMI, CIP, NOR-treated samples. (B) Venn diagram of GCSs in different FQ-treated samples. A total of 23277 potential GCSs were identified from GCS-seq. (C) Dendrogram and unsupervised hierarchical clustering of the cleavage strengths of all potential GCSs identified from MOXI, LEVO, GEMI, CIP, and NOR treatment (*n* = 3). Due to high numbers of low cleavage strength values, we used quantile break color scheme to ensure each color represents an equal proportion of the data for visualization purpose. Data were clustered using complete linkage. (D) Sequence logo representation of the central 18 nucleotides surrounding GCSs.

### Starving cultures exhibit little difference in GCS distribution from Ori-Ter

The large number of GCSs suggested that cleavage occurred extensively throughout the chromosome, and a genome atlas showing the GCS position and cleavage strength confirmed that (Figure 3A). This prompted us to examine whether GCSs from different FQ treatments exhibited any bias with respect to the macrodomains (MD) of the *E. coli* chromosome. Based on the interaction frequency of pairs of distant DNA sites using a site-specific recombination system, the *E. coli* chromosome had previously been segmented into four independent macrodomains (MDs), namely, Ori, Ter, Left, and Right and two Non-Structured (NS) regions, NS-Right and NS-Left (50). Previous studies on growing cultures have shown that GCSs are enriched in the Ori macrodomain, where DNA replication starts, whereas they are depleted in the Ter macrodomain, which is where DNA replication ends (34, 35, 61, 62). Notably, the Ori-Ter difference was observed to reach up to 4-fold (34). However, in growth-inhibited cultures, the MD bias of GCSs remained an open question. Here, for all FQ treatments the GCS strength in the Ori region was approximately equal to that expected for a domain of that size if GCSs were distributed randomly (Figure 3B). Ter showed a significantly lower GCS strength than the expected value for 1 out of the 5 of FQs (MOXI). When we performed the analysis based on the number of distinct GCSs, rather than cumulative strength, Ori and Ter were either near their expected levels or slightly below depending on the FQ used (Supplementary Figure S8). Overall, our data suggests little MD bias with respect to GCSs in stationary-phase cultures.

**Figure 3:**
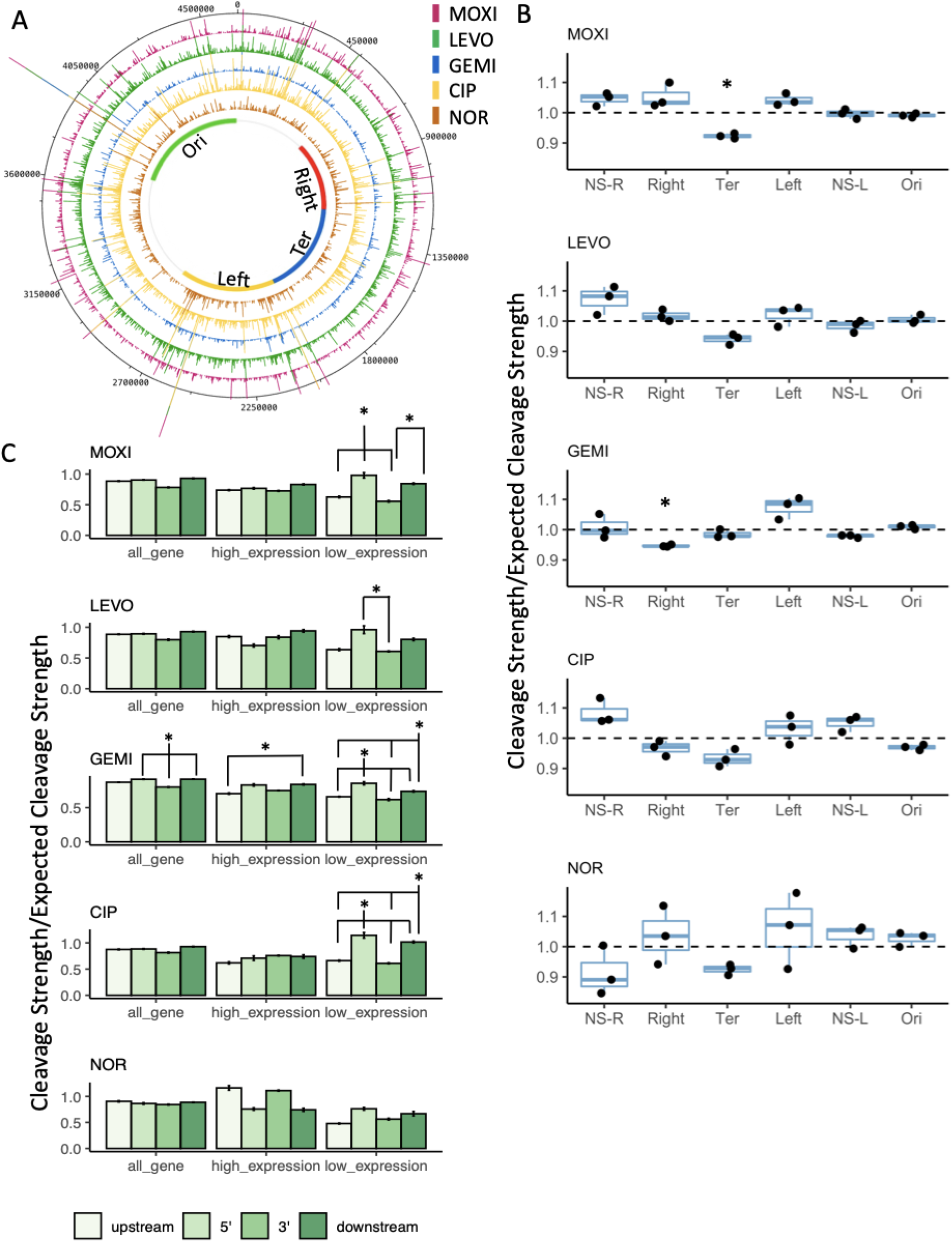
GCSs distributed across the chromosome do not show Ori-Ter gradient nor enrichment downstream of highly transcribed genes. (A) Genome atlas representation of GCSs location distribution and cleavage strengths. Inner circle depicts the macrodomain (MD) boundaries (Supplementary Table S3). (B) Cumulative gyrase cleavage strength within each MD were measured and normalized with respect to the length of the MD following MOXI, LEVO, GEMI, CIP, and NOR treatment. A value of 1.0 indicates that the cumulative cleavage strength equals the expected cumulative cleavage strength. Data are presented as box plots showing the individual normalized cleavage strength (*n* = 3). Statistical analysis was performed using one-sample t-test comparing to value of 1. Asterisk (*) denotes statistical significance (adjusted *p* < 0.05). (C) Bar plot representation of the normalized gyrase cleavage strength in upstream, 5’, 3’ and downstream of all genes, genes with high expression, and genes with low expression following FQ treatments. Statistical analysis was performed using one-way ANOVA assessing the effects of location to cleavage strength (LEVO all genes: F(3,8) = 0.46, p = 0.72; LEVO high expression genes: F(3,8) = 1.39, p = 0.31; LEVO low expression genes: F(3,8) = 4.9, p = 0.03; MOXI all genes: F(3,8) = 1.87, p = 0.21; MOXI high expression genes: F(3,8) = 1.01, p = 0.44; MOXI low expression genes: F(3,8) = 15.1, p < 0.002; GEMI all genes: F(3,8) = 8.76, p = 0.007; GEMI high expression genes: F(3,8) = 0.004, p = 0.44; GEMI low expression genes: F(3,8) = 46.82, p =2.03e-5;CIP all genes: F(3,8) = 0.34, p = 0.797; CIP high expression genes: F(3,8) = 0.004, p = 0.44; CIP low expression genes: F(3,8) = 16.16, p =0.0009;NOR all genes: F(3,8) = 0.031, p = 0.992; NOR high expression genes: F(3,8) = 1.387 p = 0.315; NOR low expression genes: F(3,8) = 1.098, p =0.405) followed by Tukey HSD post hoc test for multiple comparisons. Asterisk (*) denotes statistical significance (adjusted *p* < 0.05) between indicated regions.

To investigate possible GCSs distribution biases independent of MDs, we divided the chromosome evenly into 10 bins (A through J) and conducted comparable analyses. Again, we did not find enrichment using cumulative cleavage strength or the number of distinct GCSs in bins I or J (overlaps with Ori), whereas bin D, which contains *dif*, consistently showed the lowest cumulative cleavage strength and number of distinct GCSs for all FQ treatments (Supplementary Figure S9). Despite these statistically significant depressions, the fold differences in the distinct number of GCSs between Ori and Ter were modest and much smaller in the growth-inhibited cultures studied here compared to what was observed for growing cultures previously (34). For instance, the maximum fold difference in number of distinct GCSs found in bin I, where *oriC* lies, versus those found in bin D, where *dif* resides, was 1.19 (NOR), with a mean fold difference of 1.11 across all FQs tested, compared to > 4-fold differences for the same comparison in actively-growing populations (34, 63). Together, these data suggest extensive DNA gyrase activity along the chromosome in stationary-phase cultures that shows little Ori-Ter gradient.

### GCS distributions do not show significant associations with transcriptional activity

Past studies have shown that GCSs are influenced by transcription-induced supercoils in growing cultures where enrichment was observed downstream of highly-transcribed genes (61, 64). To assess the association between transcription and cleavage activity in starved cultures treated with different FQs, we measured the gene expression of stationary-phase cultures with RNA-seq prior to the addition of FQs. The associated cumulative cleavage strength found in the upstream, 5’, 3’, or downstream regions of highly transcribed genes (normalized expression mean > 400 transcripts per million (TPM), approximately top 10% of transcribed genes), genes with low levels of transcription (normalized expression mean < 1.5 TPM, approximately bottom 10% of transcribed genes), and all genes were calculated. We did not observe significant enrichment downstream of highly transcribed genes when comparing to the upstream, 5’ or 3’ regions for the majority of the FQs (except GEMI) (Figure 3C), and a similar observation was found when all genes were considered. Interestingly, for genes with low expression levels, we observed a cleavage bias towards the 5’ region, followed by downstream region, which was consistent across the 5 FQs tested (Figure 3C).

Additionally, we assessed the distribution of number of distinct GCSs with respect to transcriptional activity and again and did not find significant differences when comparing the upstream, 5’, 3’, or downstream of highly transcribed genes in 4 out of 5 FQs tested, whereas GEMI showed a slight depression of number of distinct GCSs in the upstream region. (Supplementary Figure S10). Despite this statistically significant depression, the region with the highest number of distinct GCSs (3’) only showed a 24% increase in number of distinct GCSs compared to the lowest abundance region (upstream), which was far lower in magnitude compared to what had been found in a similar analysis performed in nutrient-replete conditions (34). Specifically, the downstream region (where the highest number of distinct GCSs was found) had approximately twice as many GCSs than the upstream region in genes with high transcription levels when treated with CIP (34).

### GCS distributions are negatively correlated with FQ persister levels

Given the differences in GCS distributions we observed for distinct FQs, we sought to determine if any features of those distributions could explain variations in persister levels. The first feature we investigated was whether the number of distinct GCSs, which we reasoned should reflect the potential sites of DNA damage in a population, correlated with persister survival. To test this, we performed persistence assays with the five FQs used above on stationary-phase cultures at the same concentrations as used in GCS-seq. Assays showed a range of persister levels: for MOXI, LEVO, GEMO, CIP, and NOR, persisters were approximately 1 in 1000, 1 in 50, 1 in 20, 1 in 10, and 1 in 2, respectively, following a 5 h course of treatment (Supplementary Figure S5). Examination of persister levels as a function of distinct GCSs revealed a negative association where fewer distinct GCSs corresponded to higher survival (Figure 4A). For example, MOXI had the highest number of distinct GCSs and the lowest level of persisters, whereas NOR had the fewest GCSs and highest survival after treatment. To probe the strength of the correlation, we performed linear regression on the log-transformed survival fraction with respect to the number of distinct GCSs. The coefficient of determination (*R*^2^) was 0.83, showing that the variation in the number of GCSs could explain 83% of the variation in survival. These results suggested that the number of distinct GCSs was negatively correlated with survival and thus the more potential sites of DNA damage from a FQ translates to lower persister levels. We further investigated whether cumulative cleavage strength, which is a metric that should reflect the extent of DNA damage in a population, correlated with persistence. As depicted in Figure 4B, we found an even stronger negative correlation with *R*^2^ = 0. 98 between the log survival fraction and cumulative cleavage strength. This strong negative correlation implied that the cumulative ability of a FQ to stabilize DNA gyrase across the chromosome is highly predictive of persister levels.

**Figure 4:**
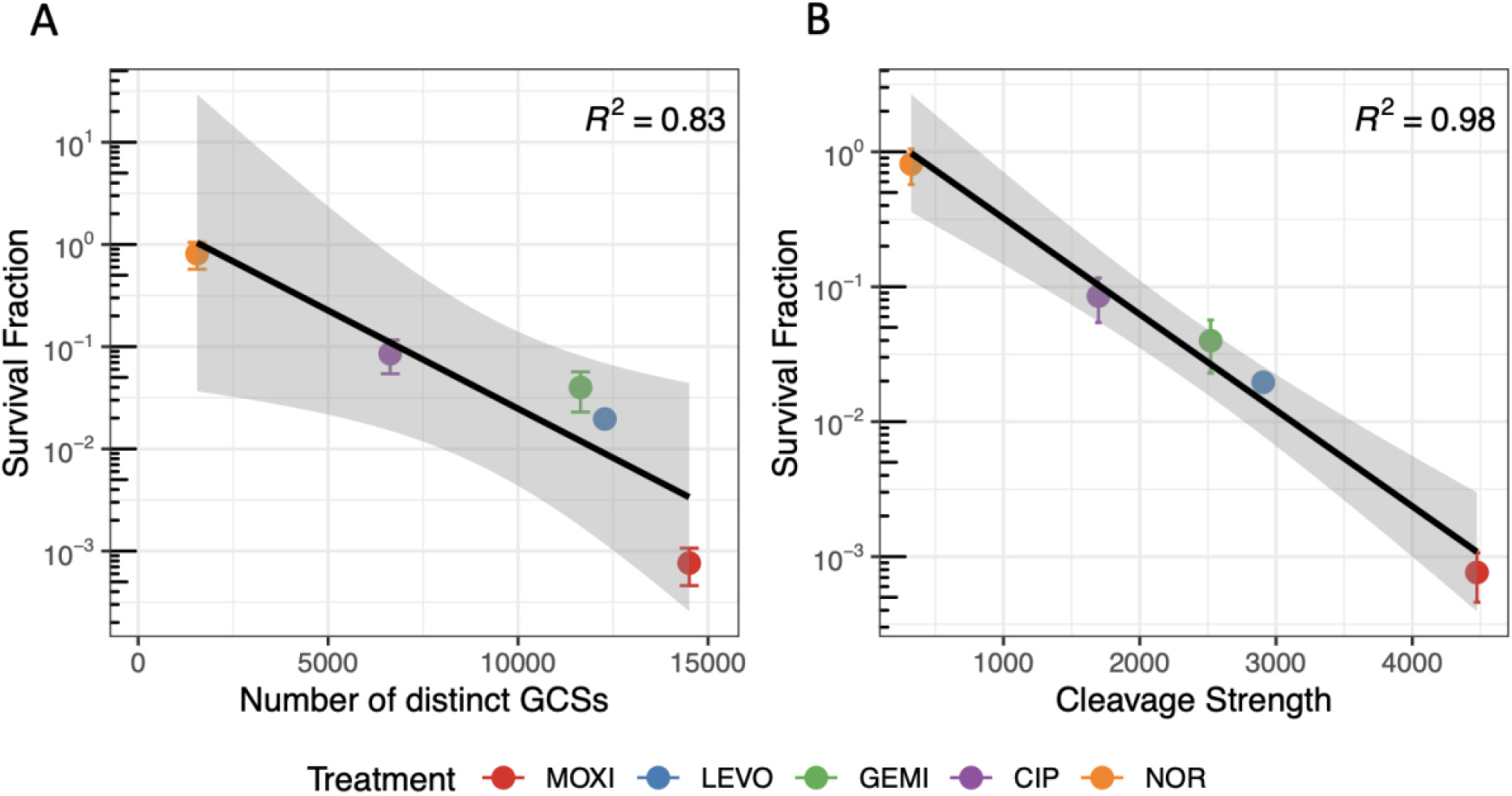
Persister survival is negatively correlated with cleavage strength and number of distinct GCSs. Stationary-phase *E. coli* cultures were treated with MOXI, LEVO, GEMI, CIP, or NOR for 5 h and the survival fractions were measured. Data are presented as mean survival fraction ± SEM (*n* = 3) against (A) the number of distinct GCSs identified for each FQ treatment, or (B) the cumulative cleavage strengths for each FQ treatment. Black curve represents the fitted regression line and coefficient of determination (*R*^2^) is indicated in each plot. Shaded bands around the regression line represent 95% confidence interval for the regression estimates.

### Modulation of individual GCSs near Ori or Ter does not impact FQ persistence

In addition to the number of distinct GCSs and cumulative cleavage strength, we considered whether the chromosomal locations of GCSs could impact persister levels. To begin, we performed persistence assays on strains Mu_origin 1 and MuScr_origin 1, which contain a strong GCS and non-cleaved control close to *oriC*, respectively. Following FQ treatment, we did not observe any statistically significant difference in persister levels between Mu_origin 1 and MuScr_origin 1 (Figure 5A-C and Supplementary Figure S11AB). To assess the generality of these results, we inserted Mu and MuScr in a different location close to *oriC* (Mu_origin 2 and MuScr_origin 2) and performed persistence assays. We again did not observe any differences in survival between Mu and MuScr strains (Figure 5A-C and Supplementary Figure S11AB). These results suggested that altering a strong GCS near *oriC*, which is where DNA replication begins, did not have observable impacts on FQ persistence.

**Figure 5.**
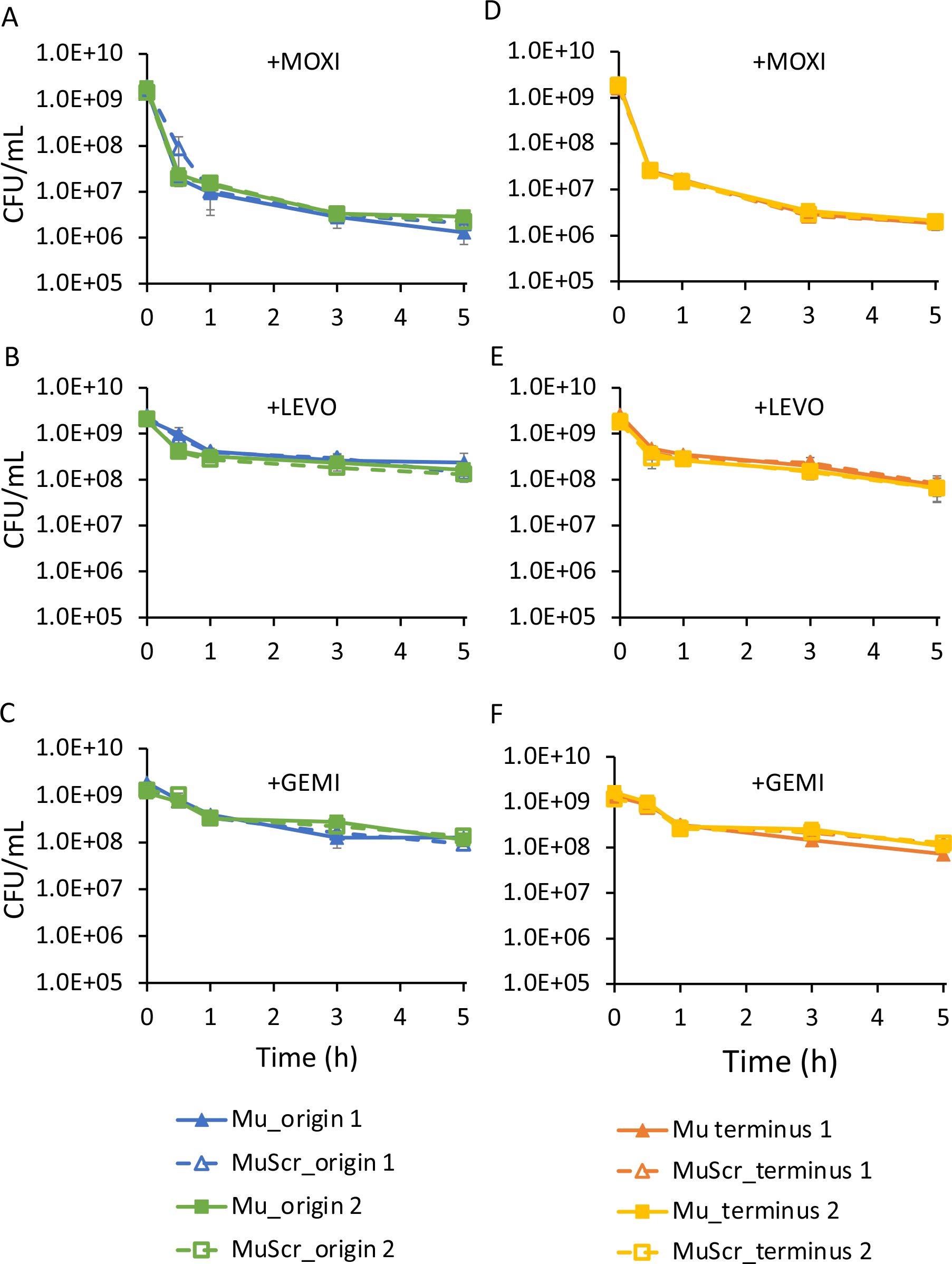
Introduction of strong GCS close to origin or terminus does not impact survival. Wild type *E. coli* strain MG1655 was modified by insertion with either a confirmed strong gyrase cleavage sequence Mu or a scrambled control sequence MuScr at positions close to the origin (A-C) or terminus (D-F). Cultures with inserted sequences were grown to stationary phase and then treated with either 5 μg/ml MOXI (A and D), LEVO (B and E), or GEMI (C and F). Additional data with CIP and NOR treatment and water treatment controls can be found in Supplementary Figure S11 and S12, respectively. Data denote means ± SEM (*n* ≥ 3). P = 0.05 (two-tailed *t*-tests with unequal variances) was used as significance threshold and log transformed CFU/mL values were compared between Mu and MuScr strains at each insert location for each time point. Statistical significance was not detected.

Another genomic location we probed was near *dif*, which is close to the termination site of DNA replication (65). Analogously, we introduced strong GCSs and their scrambled equivalents at two distinct sites near *dif*. Persistence assays again showed comparable survival between strains with an additional strong GCS or its scrambled equivalent (Figure 5D-E and Supplementary Figure S11CD). The water-treated controls demonstrated that killing required the addition of FQ (Supplementary Figure S12), and that all Mu sites considered here were confirmed to be strong GCS (Supplementary Figure S13).

### Removal of a strong GCS near *recB* did not impact FQ persistence

Since incorporating a strong GCS had no observable impacts on FQ survival, we investigated whether removing a strong GCS near an important DNA repair gene would impact persister levels. Among the 956 shared GCSs, a top-ranked site according to cleavage strength (No. 11 in LEVO, No. 3 in MOXI, No. 9 in NOR, No. 4 in CIP, and No. 12 in GEMI) was found in the *recD* opening reading frame (ORF) (66). *recB* and *recC* have been found to be important to FQ persistence previously (21, 31, 67), and *recD* neighbors *recB* in the same operon (68). To remove the GCS in *recD*, we began by deleting the entire gene, rationalizing that if an impact were observed, we would then more precisely modify the GCS in follow-up experiments. However, we found that the knockout mutant exhibited indistinguishable survival compared to wild-type for all the FQs tested (Figure 6). These data demonstrated that removal of a strong GCS, even one adjacent to a critical DNA repair gene (*recB*), did not impact persister levels.

**Figure 6:**
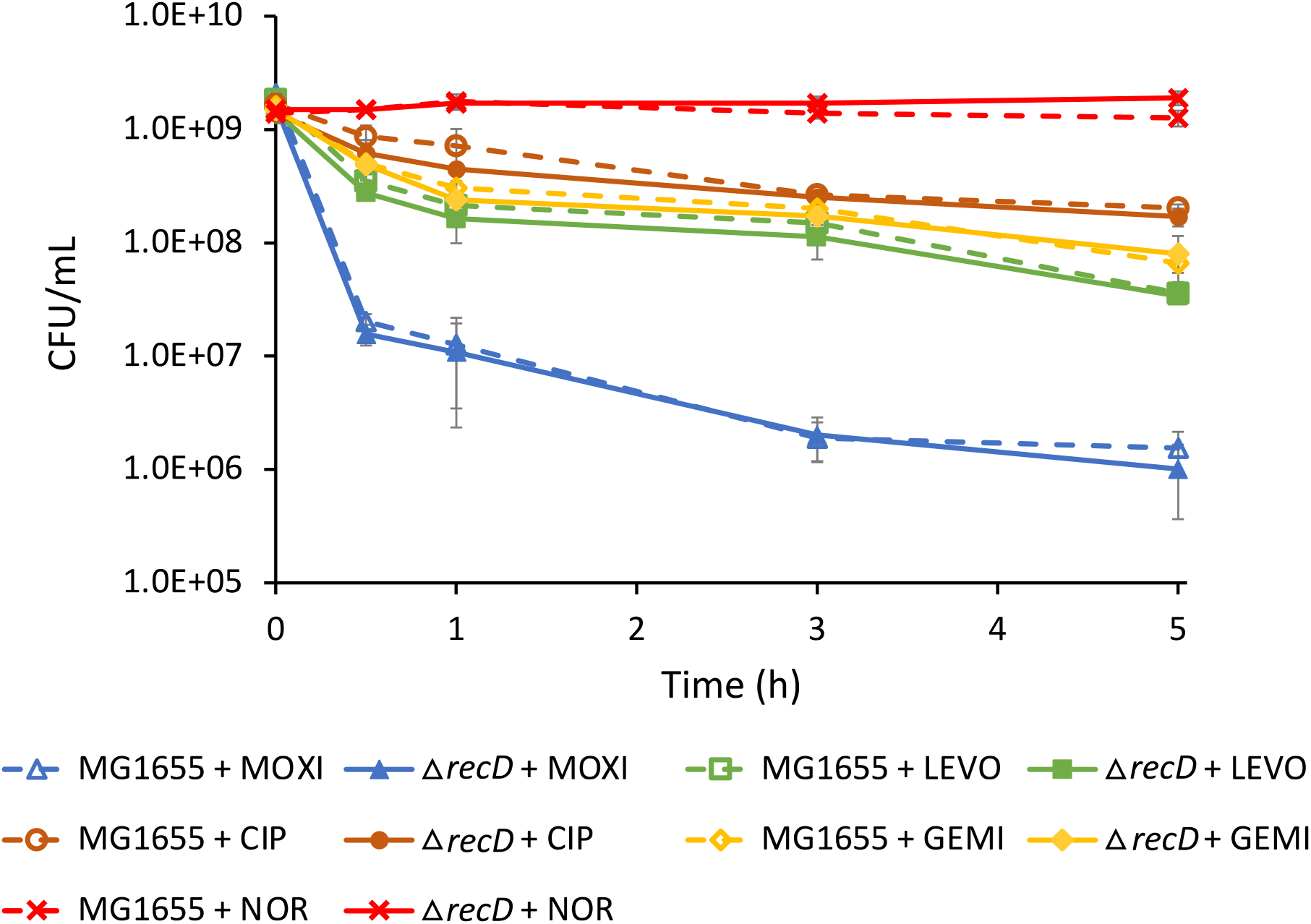
Removal of a strong GCS within *recD* does not impact survival. Wild type *E. coli* strain MG1655 and Δ*recD* were grown to stationary phase and treated with MOXI, LEVO, GEMI, CIP, or NOR. Data denote means ± SEM (*n* ≥ 3). P = 0.05 (two-tailed *t*-tests with unequal variances) was used as the significance threshold, comparing log transformed CFU/mL values between MG1655 and Δ*recD* for each FQ treatment. Statistical significance was not detected.

## Discussion

Persisters are enriched in biofilms and granulomas that protect bacteria from immunity (11, 13), and have been associated with chronic infections such as tuberculosis, cystic fibrosis, and urinary tract infections (5, 11, 69–73). Recent works have also shown that persisters can facilitate the development of resistance (8, 10, 74) and that just a single round of FQ treatment was sufficient to enhance resistance development significantly (8). Alarmingly, the frequency of persisters in a susceptible population (*e.g*., FQ:1×10^−2^ to 1×10^−5^ (21, 31, 75)) far exceeds that of spontaneously resistant mutants (FQ: 1×10^−7^ or less (76)), which suggests that persisters have a higher likelihood of causing initial treatment failure. For these reasons, a better understanding of persister physiology and survival mechanisms could improve treatment outcomes for relapsing and chronic infections.

Persisters had previously been thought to survive treatment due to a lack of antibiotic-induced damage (4, 77–79). However, that model does not apply universally, and FQs are a prime example of persisters surviving despite antibiotic-induced damage (21, 31–33, 74). Prompted by the capacity of FQs to interact with their primary target in persisters and non-persisters alike (21), we investigated whether characteristics of that interaction would be impactful on persistence. Here, we measured the genome-wide GCS distributions after treatment with five FQs in stationary-phase *E. coli* cultures using GCS-seq. We identified a total of over 20000 cleavage sites along the *E. coli* genome, which is equivalent to approximately 1 in every 230 nucleotides. Such extensive cleavage had been noted before for growing cultures (34), but was somewhat surprising in the ATP-depleted stationary-phase cultures considered here (80). Location and cleavage strength depended on the type of the FQ used, and notably, cumulative cleavage strength was found to be an extremely strong predictive variable for persister levels (*R*^2^ = 0.98). Further, modulation of individual GCSs failed to alter persister levels, which together suggested that survival is likely to be a function of the cumulative damage across the genome rather than damage to specific loci. Therefore, we postulate that FQs that better stabilize DNA gyrase in cleaved complexes will produce lower persister levels.

Beyond persistence, the data generated revealed differences and similarities with respect to GCSs between growing and non-growing cultures. Previous work has shown GCS enrichments near Ori and depletions near Ter due to active replication and transcription in nutritive environments (34, 63). Here, we observed little differences in the cumulative cleavage strength or the number of distinct GCSs within different MDs. We expect that this originates from the lack of DNA replication and highly reduced transcription rates (81) that are characteristic of stationary-phase cultures. Further, we did not find that GCSs were preferentially found downstream of highly transcribed genes compared to upstream in non-growing cultures, as was previously observed with growing populations (34). However, consistent with their growing counterparts, non-growing bacteria contained GCSs with a similar cleavage motif where gyrase preferentially cleaves before guanine residues (34).

In conclusion, we measured the genome-wide cleavage pattern of DNA gyrase when stabilized by different FQs in stationary phase cultures. Those GCS distributions differed from those in exponential-phase cultures with respect to location and strength, but similarly reflected a catalytic preference for DNA gyrase to cleave DNA adjacent to guanine residues. Importantly, the data presented suggests that the genome-wide GCS distribution is highly predictive of persister levels, despite such measurements being conducted on a population level rather than the persister subpopulation. We expect that the capacity of FQs to stabilize DNA gyrase in persisters and cells that die from treatment does not differ appreciably, which is why such a metric is such a predictive variable between different FQs. Taken together, these data suggest that identifying new FQs or adjuvants that better stabilize DNA gyrase in cleaved complexes will reduce persistence and potentially improve the treatment of recalcitrant infections.

## Data Availability

The main data supporting the findings of this study are available within the article and in the Supplementary Information. The code and data for analysis can be found on GitHub at https://github.com/CathyTJC/GCS-seq. Raw sequencing data were deposited with GEO under accession number GSE206610 (https://www.ncbi.nlm.nih.gov/geo/query/acc.cgi?acc=GSE206610, token: qbmriikodpwtfsv).

## Supplementary Data

Supplementary Data are available at NAR online.

## Acknowledgements

We thank Wei Wang, Jessica B. Wiggins, Robert W. Leach, Lance R. Parsons (High Throughput Sequencing Facility, Lewis-Sigler Institute for Integrative Genomics), and Saw Kyin (Princeton Proteomics and Mass Spectrometry Core Facility) for their assistance and suggestions. We also acknowledge the National BioResource Project (NIG, Japan) for distribution of the Keio Collection.

## Funding

This work was supported by the NIAID of the National Institutes of Health (R01AI130293: M.P.B.). The content is solely the responsibility of the authors, does not necessarily represent the official views of the funding agency, and the funder had no role in the design or implementation of the experiments or the decision to publish.

## Conflict of interest statement

None declared.

## Notes

### Competing Interest Statement

The authors have declared no competing interest.

