## Supplemental figures for "Genome-wide mapping of fluoroquinolone-stabilized DNA gyrase cleavage sites displays drug specific effects that correlate with bacterial persistence"

**Title**

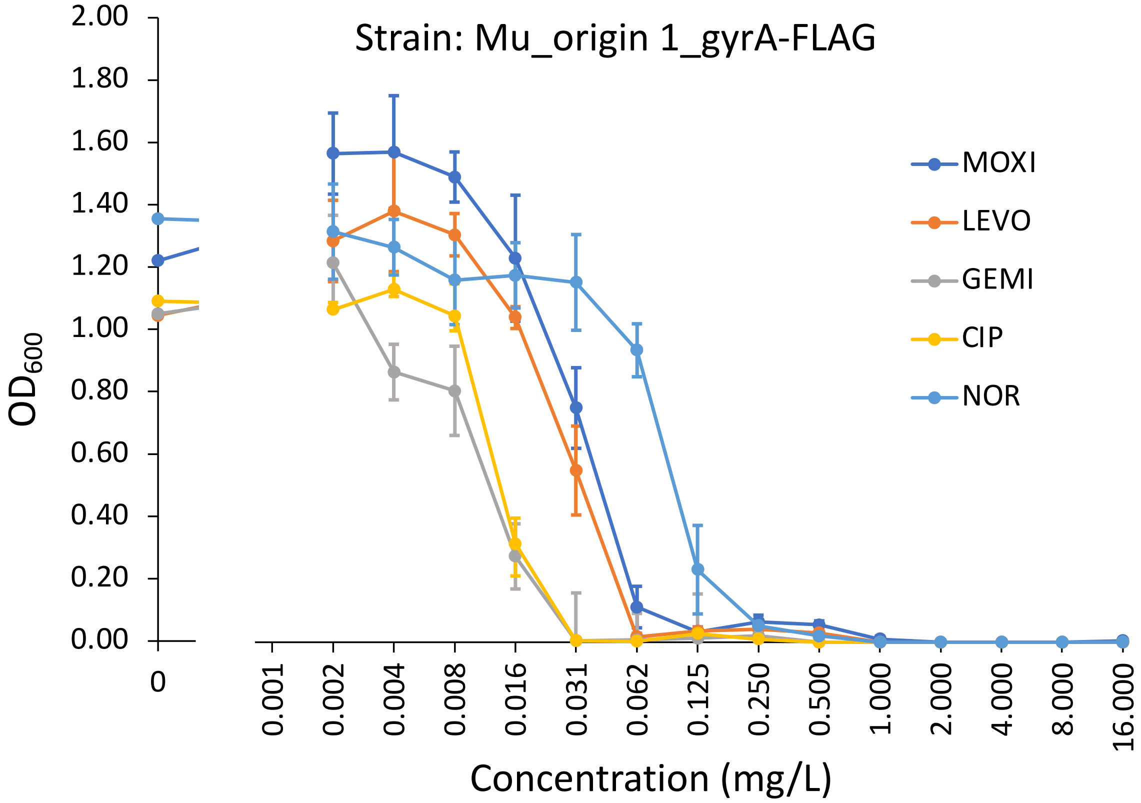


| MIC (mg/L) | MOXI | LEVO | GEMI | CIP | NOR |
|---|---|---|---|---|---|
| | 0.125 | 0.0625 | 0.03125 | 0.03125 | 0.25 |

**Supplementary Figure S1: Minimum inhibitory concentrations of Mu_origin 1_gyrA-FLAG strain.** Minimum inhibitory concentrations (MICs) were determined by microdilution method (Materials and Methods). Optical density ($OD_{600}$) was measured following 16-18 h incubation with MOXI, LEVO, GEMI, CIP, or NOR with indicated concentrations. Data represent mean $OD_{600} \pm$ SEM ($n = 6$). MICs are shown in the accompanying table.

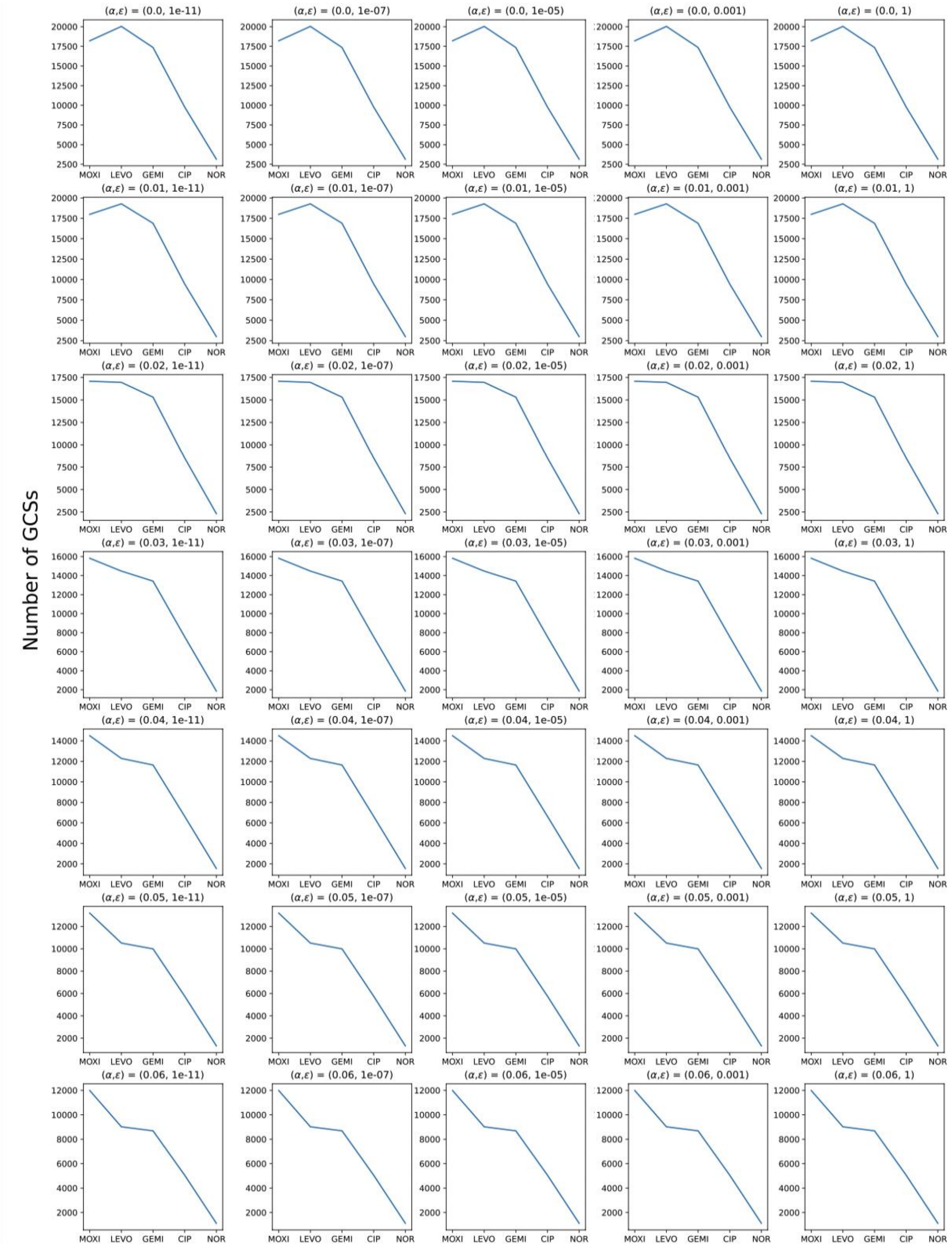


**Supplementary Figure S2: GCS-seq performance does not depend on the selection of**

**pseudo-count.** The number of GCSs identified from different FQ treatment is not affected by the choice of $\boldsymbol{\varepsilon}$ (1e-11 to 1) over varying $\boldsymbol{\alpha}$.

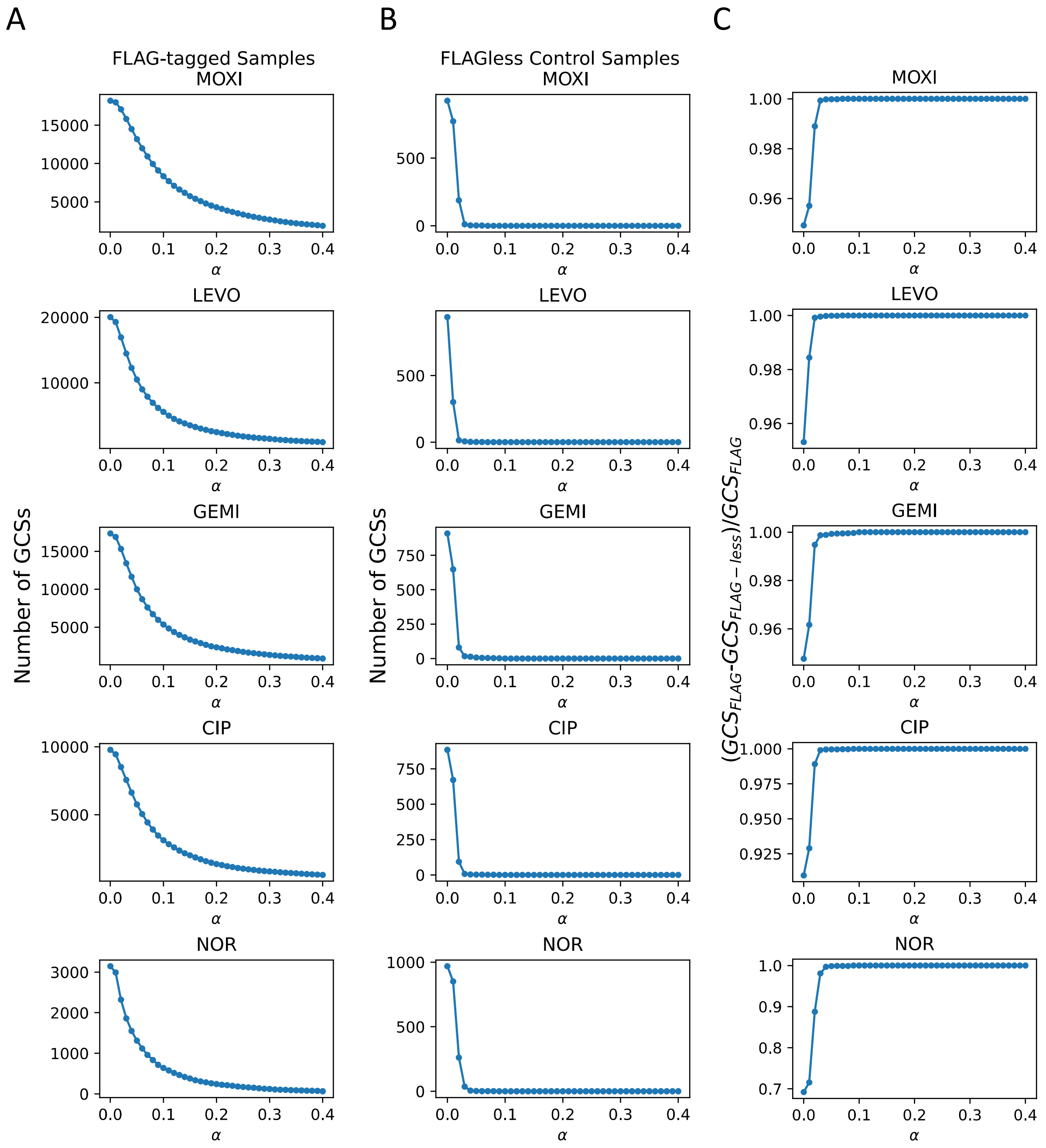


**Supplementary Figure S3: GCS calling achieves high precision.** The number of distinct GCSs identified from MOXI, LEVO, GEMI, CIP, and NOR treatment is plotted against the hyperparameter $\boldsymbol{\alpha}$ in (A) FLAG-tagged strain and (B) FLAG-less control strain. (C) The precision of the algorithm is approximated by calculating the ratio of genuine GCSs ($GCS_{FLAG}$ – $GCS_{FLAG\text{-}less}$)/$GCS_{FLAG}$). Accuracy exceeds 99% when $\boldsymbol{\alpha} \geq 0.04$.

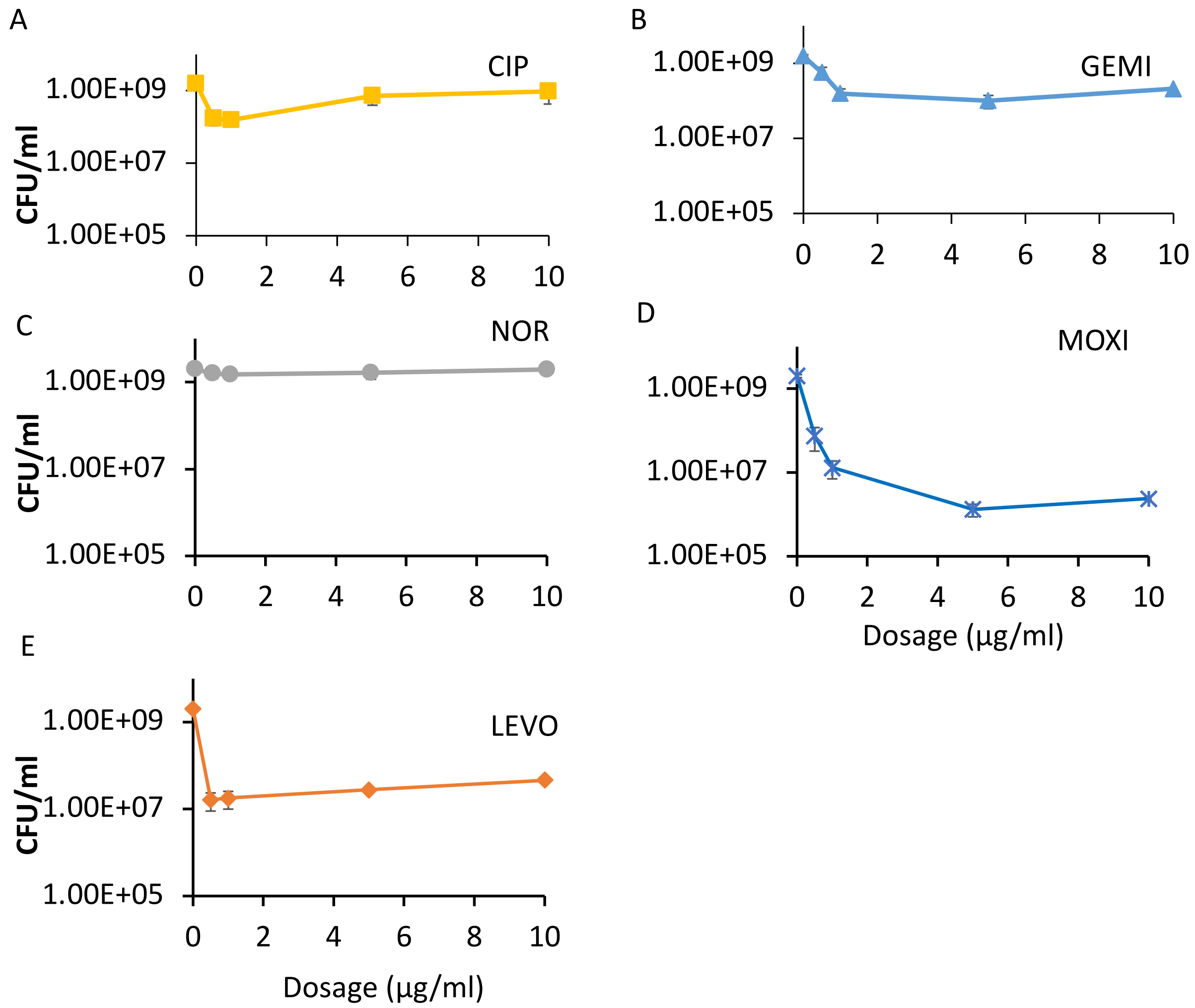


**Supplementary Figure S4: Survival of stationary-phase *E. coli* treated with different concentrations of FQs.** To determine the FQ concentration used for GCS-seq and persistence assay, cells were grown to stationary phase and treated with 0 - 10 µg/mL of (A) CIP, (B) GEMI, (C) NOR, (D) MOXI, and (E) LEVO. CFU/mL values were determined after a 5 h course of treatment.

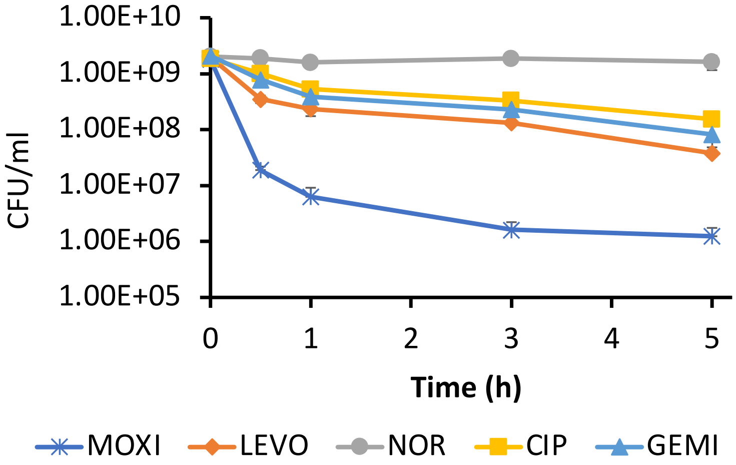


**Supplementary Figure S5: Survival of stationary-phase *E. coli* following treatment with different FQs.** Persistence assays were performed with 5 µg/mL LEVO, 5 µg/mL MOXI, 1 µg/mL CIP, 5 µg/mL GEMI, and 5 µg/mL NOR on stationary-phase Mu_origin 1_gyrA-FLAG over a course of 5 h treatment. CFUs were enumerated right before drug treatment (time 0), and at 0.5, 1, 3, 5 h following drug treatment. Data was represented as mean ± SEM ($n = 3$).

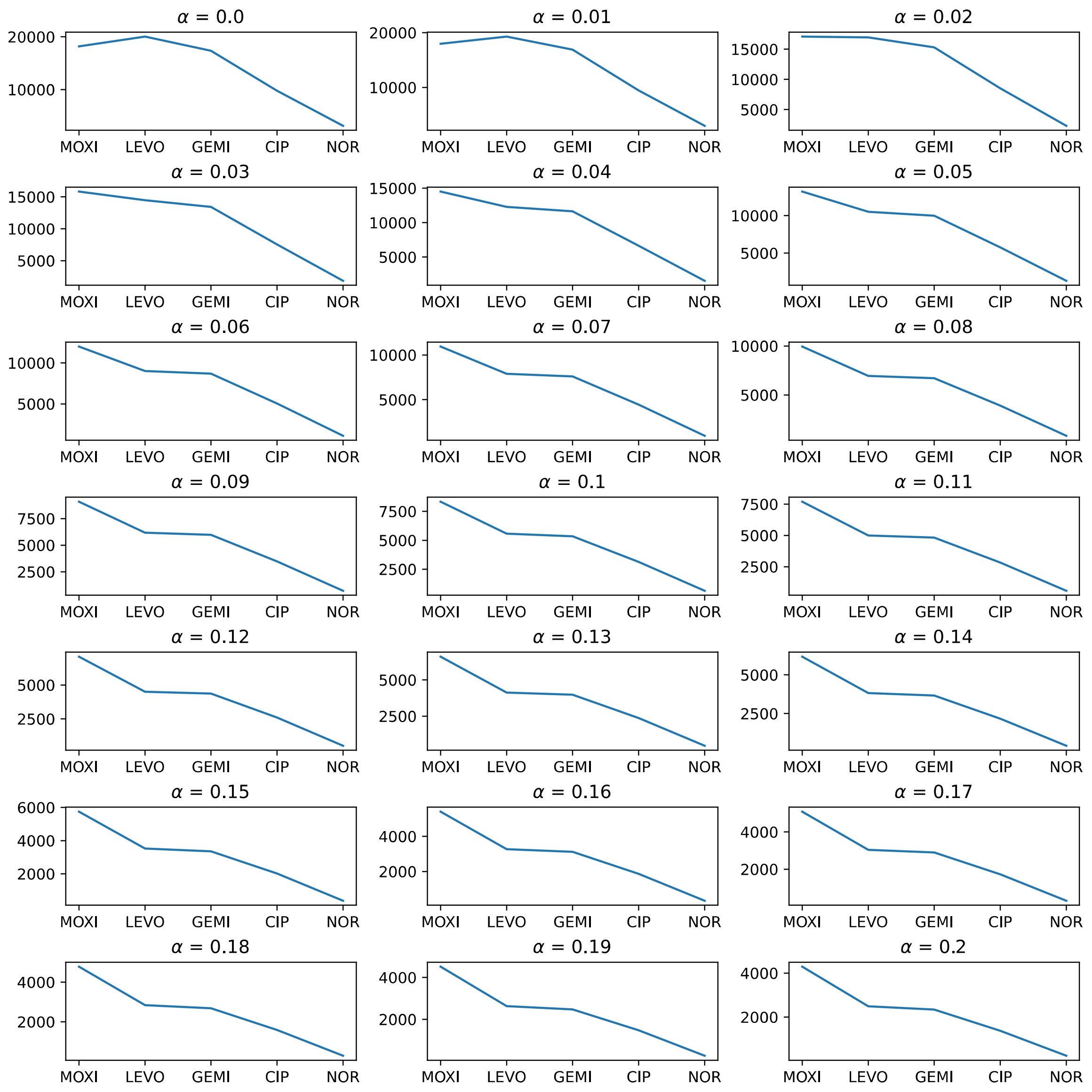

**Supplementary Figure S6: GCS calling yields a consistent order in the number of distinct GCSs with respect to FQ treatment.** The data represents the number of GCSs identified from the MOXI, LEVO, GEMI, CIP, or NOR treatment at a particular $\boldsymbol{\alpha}$ (ranges from 0.0 to 0.2). A consistent order of number of GCSs (MOXI > LEVO > GEMI > CIP > NOR) were obtained over a wide range of $\boldsymbol{\alpha}$ (0.03 to 0.20).

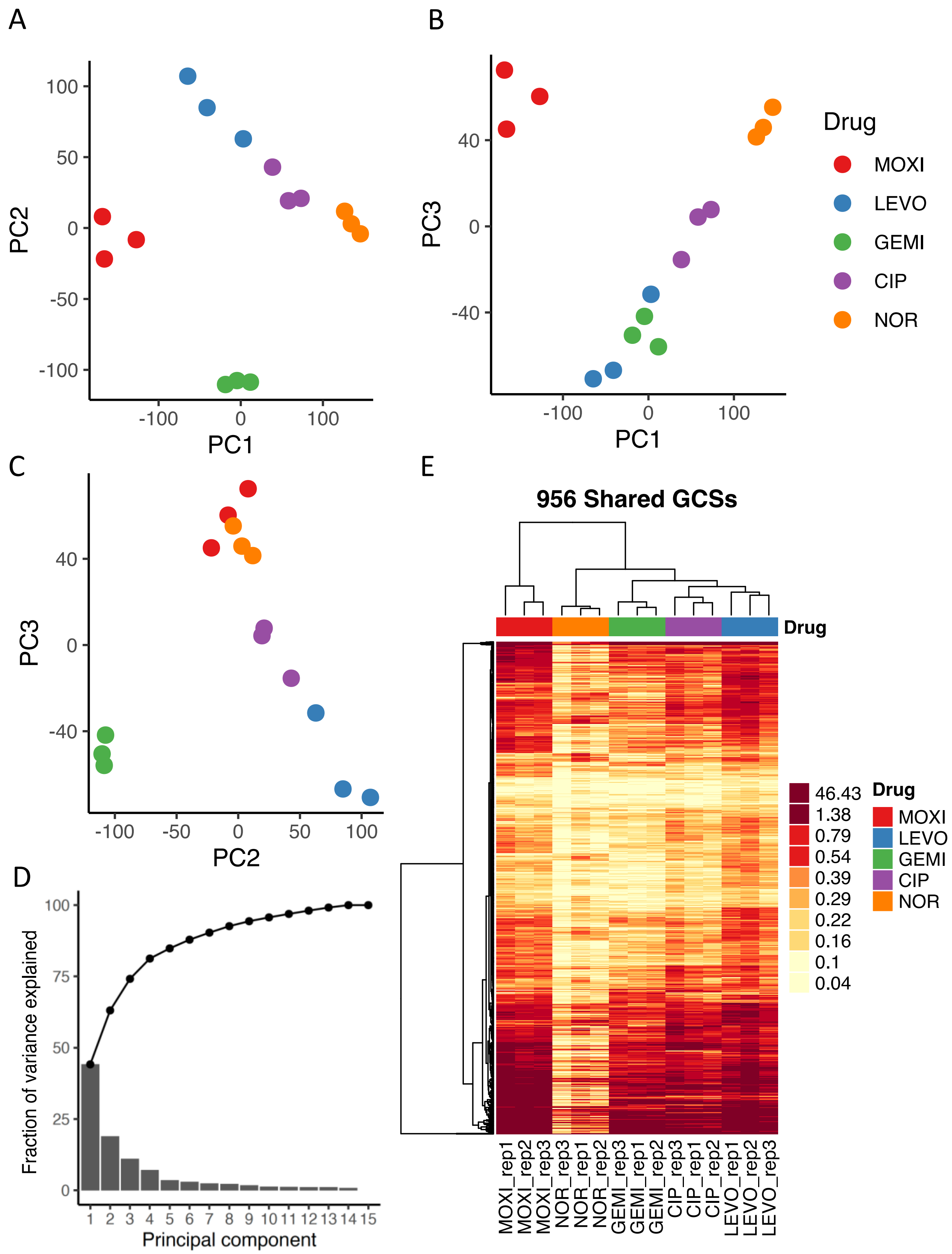

A
B
C
D
E
PC1
PC2
PC3
Drug
MOXI
LEVO
GEMI
CIP
NOR
Fraction of variance explained
Principal component
956 Shared GCSs
46.43
1.38
0.79
0.54
0.39
0.29
0.22
0.16
0.1
0.04
MOXI_rep1
MOXI_rep2
MOXI_rep3
NOR_rep3
NOR_rep1
NOR_rep2
GEMI_rep3
GEMI_rep1
GEMI_rep2
CIP_rep3
CIP_rep1
CIP_rep2
LEVO_rep1
LEVO_rep2
LEVO_rep3

**Supplementary Figure S7: GCS distributions show drug specific cleavage patterns**. (A-C) PCA analysis of gyrase cleavage strength dataset. PC1-PC2-PC3 score plots show clustering in biological replicates treated with the same FQ. (D) Screen plot of fraction of variance explained by principal components. The first 3 PCs explained ~75% of the variance. (E) Dendrogram and unsupervised clustering heatmap representation of 956 shared GCS cleavage strengths ( $n = 3$). Samples were clustered based on complete linkage. A quantile color scheme was applied for visualization purpose such that each color represents same proportion of data. The data show clustering of samples according to FQ type as well unique cleavage patterns for each FQ treatment.

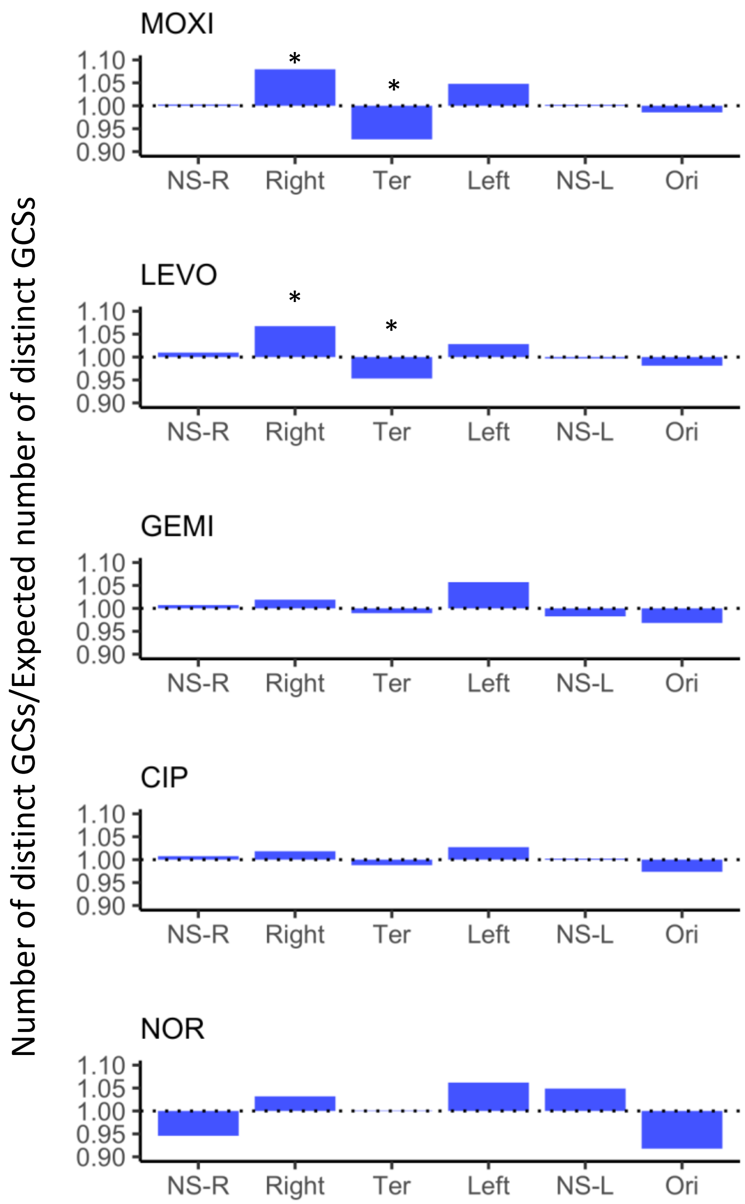


**Supplementary Figure S8: GCS distributions across MDs.** The number of distinct GCSs found with each MD were counted following MOXI, LEVO, GEMI, CIP, and NOR treatment. For visualization purpose, the number was normalized with respect to the expected number of GCSs found within each MD according to the MD lengths. A value of 1.0 indicates that the actual number of GCSs equals the expected number of GCSs. Exact binomial test was performed to determine MD effects on the number of GCSs. Asterisk ($*$) denotes statistical significance (adjusted $p < 0.05$).

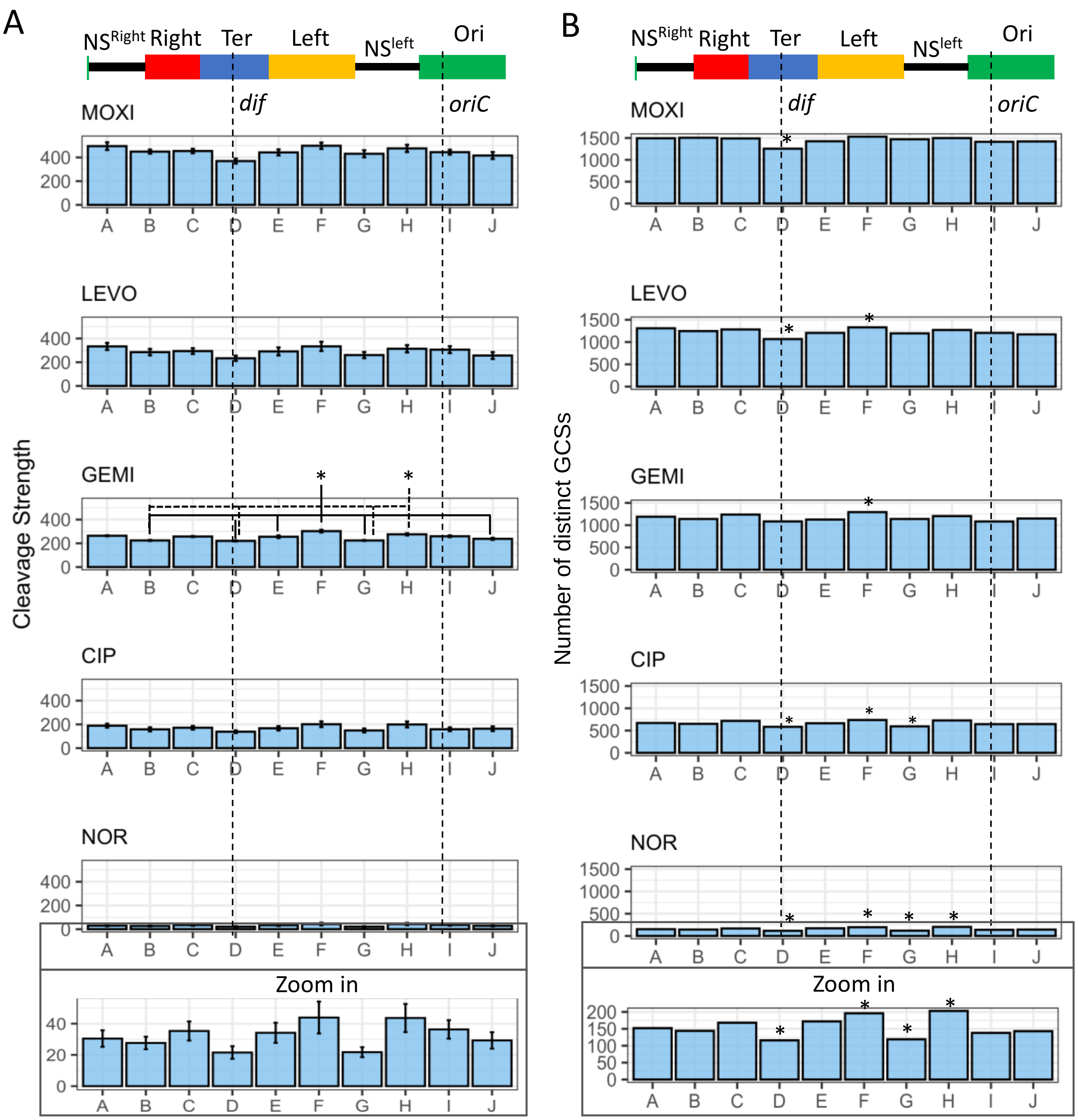


**Supplementary Figure S9: Distribution of FQ-mediated GCSs across genome without binning on MDs.** The chromosome was divided into 10 bins and (A) the cumulative cleavage strength (mean ± SEM, $n = 3$) or (B) the number of GCSs are represented. The MDs boundary are marked at top. For (A), statistical analysis was performed using one-way ANOVA assessing the effects of location to cleavage strength (LEVO: $F(9,20) = 1.28$, $p = 0.31$; MOXI: $F(9,20) =$

2.26, p = 0.06; GEMI: F(9,20) = 13.45, p = 1.11e-06; CIP: F(9,20) = 1.351, p = 0.27; NOR: F(9,20) = 1.544, p = 0.2) followed by Tukey HSD post hoc test for multiple comparisons. Asterisk ($*$) denotes statistical significance (adjusted $p < 0.05$) between indicated regions. For (B), statistical analysis was performed by exact binomial tests. Asterisk ($*$) denotes statistical significance (adjusted $p < 0.05$).

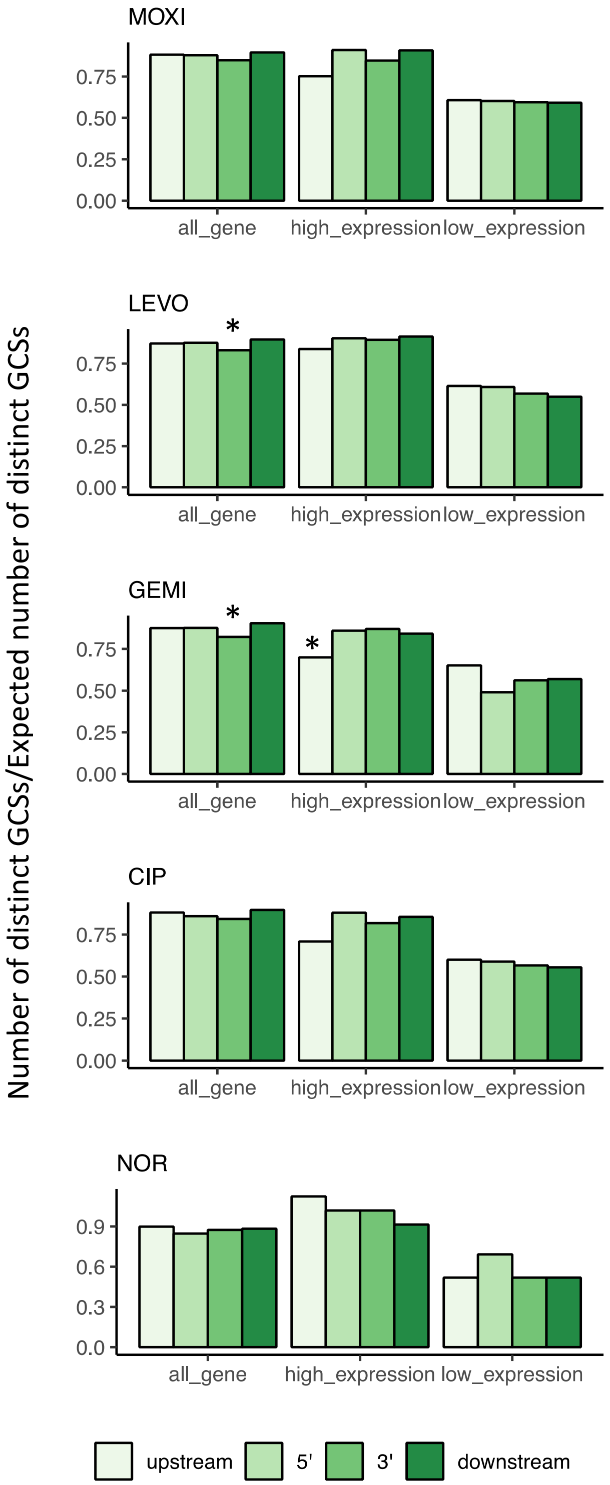

MOXI
LEVO
GEMI
CIP
NOR
Number of distinct GCSs/Expected number of distinct GCSs
0.00
0.25
0.50
0.75
0.0
0.3
0.6
0.9
*
all_gene
high_expression
low_expression
upstream
5'
3'
downstream

**Supplementary Figure S10: GCSs show no enrichment downstream highly transcribed regions.** As in Figure 3C, the genes were grouped into 3 categories: (i) all genes, (ii) genes with high expression, and (iii) genes with low expression following FQ treatment. The number of GCSs found in the upstream, 5', 3' and downstream of each category were counted. Bar plots represent the normalized number of GCSs with respect to the lengths of each category. Statistical analysis was performed by exact binomial tests assuming the unbiased probability be 0.25 (each category owns 4 regions of same length). Asterisk ($*$) denotes statistical significance (adjusted $p < 0.05$).

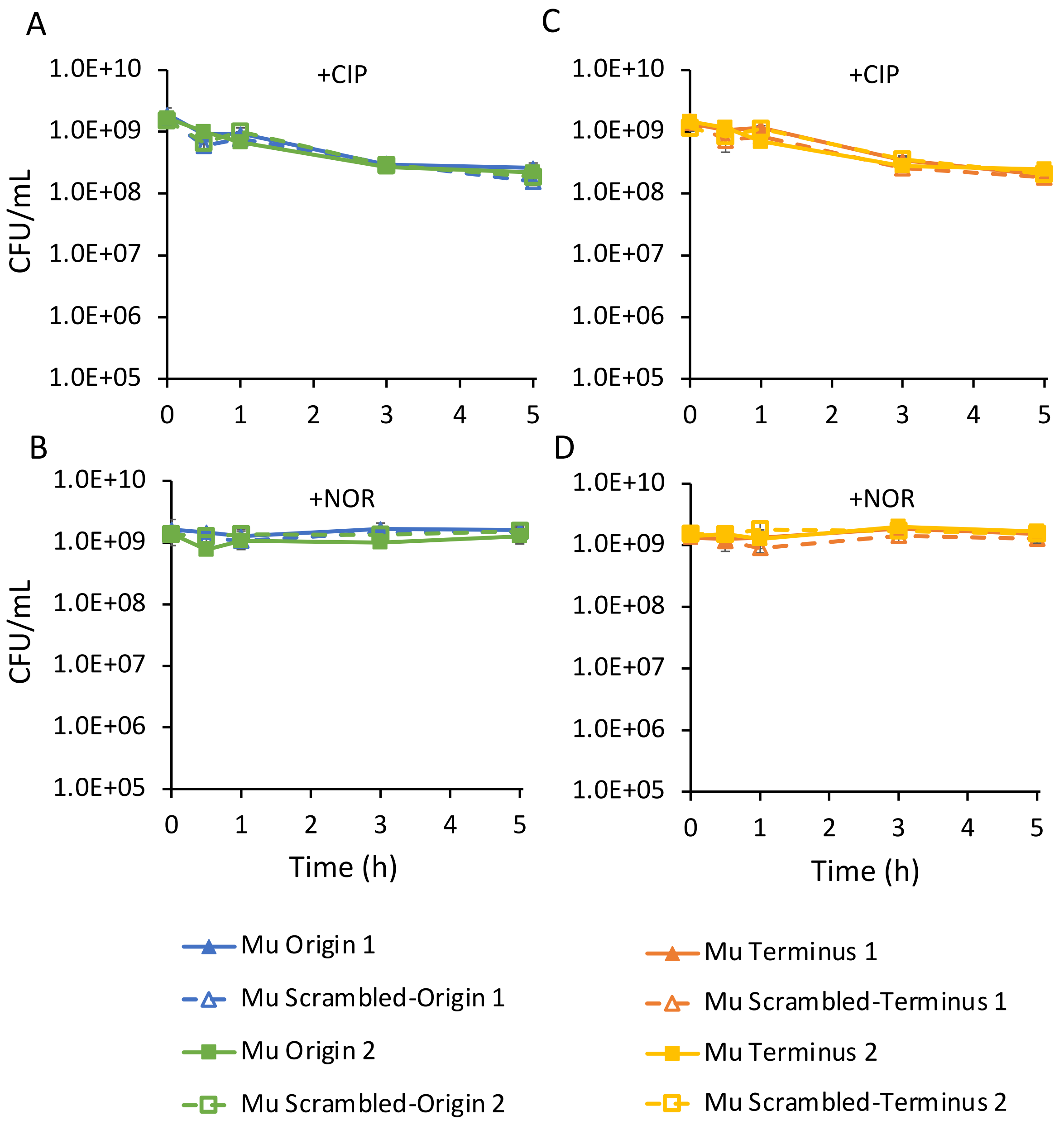


**Supplementary Figure S11: Introduction of strong GCS close to origin or terminus does not impact survival.** Data complements that in Figure 5. Analogous experiments as Figure 5 except with 1 µg/mL CIP (A and C) or 5 µg/mL NOR (B and D) treatment. Data denotes means ± SEM ($n \geq 3$). P = 0.05 (two-tailed *t*-tests with unequal variances) was used as significance threshold and log transformed CFU/mL values were compared between Mu and MuScr strains at each insert location. Statistical significance was not detected.

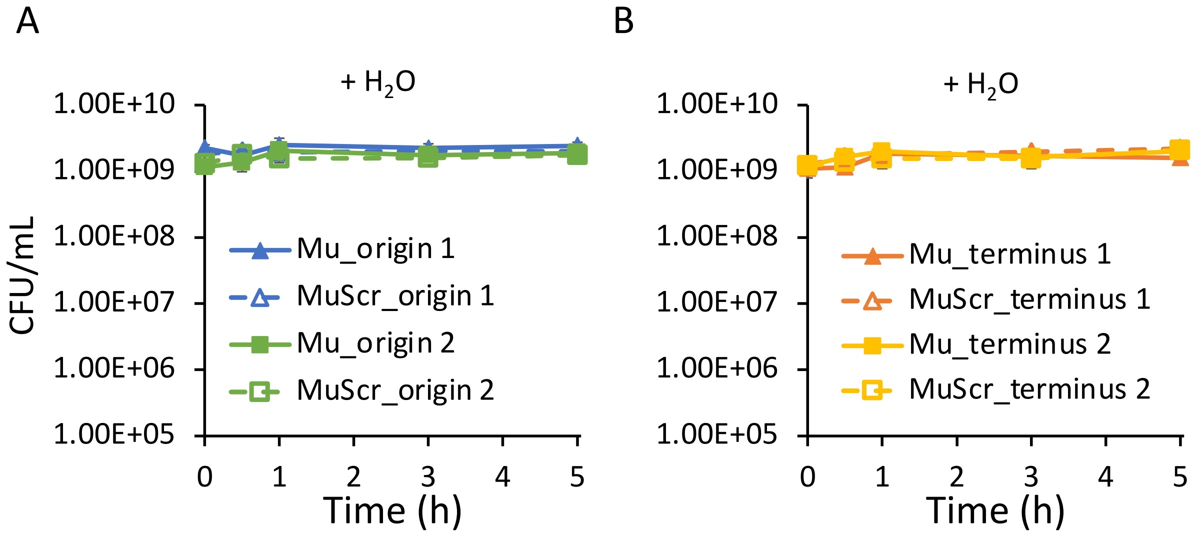


**Supplementary Figure S12: Water treatment controls for persistence assays.** Cells were grown to stationary phase and treated with water. Samples were taken at designated time points and CFU/mL values were determined. Data denotes mean ± SEM ($n = 3$). P = 0.05 (two-tailed *t*-tests with unequal variances) was used as significance threshold and log transformed CFU/mL values were compared between Mu and MuScr strains at each insert location. Statistical significance was not detected.

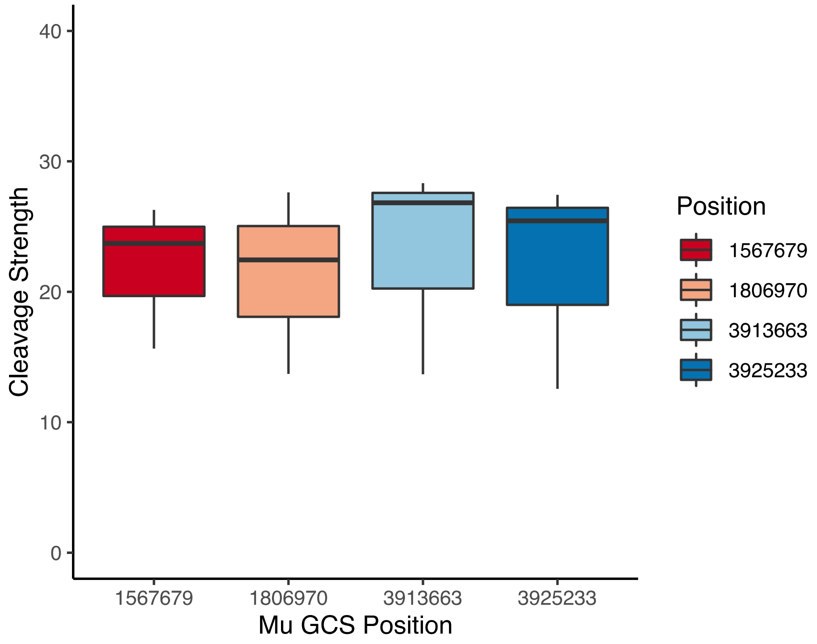


**Supplementary Figure S13: Validation of Mu cleavage when inserted close to origin or terminus.** GCS-seq was performed on strain MG1655 4Mu_gyrA-FLAG where Mu sequence was inserted to different locations (2 close to origin and 2 near the terminus, same insertion locations as shown in Figure 5 and Supplementary Figure S11). The GCSs and their associated cleavage strengths were determined following the GCS calling procedure. Data represent box plot of the mean cleavage strength at each Mu site ($n = 3$). All four Mu sites were identified as strong gyrase cleavage sites.
