## Supplemental tables for "Genome-wide mapping of fluoroquinolone-stabilized DNA gyrase cleavage sites displays drug specific effects that correlate with bacterial persistence"

**Supplementary Table S1. Bacterial strains and plasmids**

| **Strain** | **Relevant genotype** | **Source or Reference** |
|---|---|---|
| MG1655 | F-, λ-, *ilvG-*, *rfb-50*, *rph-1* | ATCC 700926 (1) |
| Mu_origin 1 | MG1655 *pstS-glmS*::*cat-Mu* | This work; Integration of 1241bp of *cat*-Mu in the intergenic region between *pstS* and *glmS* using the method of Datsenko and Wanner (2) |
| Mu_origin 1_gyrA-FLAG | Mu_origin 1 *gyrA-FLAG-kan* | This work; Integration of FLAG-*kan* at C-terminus of *gyrA* using Datsenko and Wanner method (2) |
| Mu_origin 1_gyrA-FLAGless | Mu_ origin 1 *gyrA-kan* | This work; Integration of *kan* at C-terminus of *gyrA* using Datsenko and Wanner method (2) |
| Mu_origin 2 | MG1655 *aptI-rsmG*::*Mu* | This work; Integration of Mu-*kan* in the intergenic region between *aptI* and *rsmG* using Datsenko and Wanner method and cured of *kan* using pCP20 (2) |
| Mu_terminus 1 | MG1655 *yddY-yddW*::*Mu* | This work; Integration of Mu-*kan* in the intergenic region between *yddY* and *yddW* using Datsenko and Wanner method and cured of *kan* using pCP20(2) |
| Mu_terminus 2 | MG1655 *ydiY-pfkB*::*kan-Mu* | This work; Integration of Mu-*kan* in the intergenic region between *ydiY* and *pfkB* using Datsenko and Wanner method (2) |
| MuScr_origin 1 | MG1655 *pstS-glmS*::*cat-MuScr* | This work; Integration of 1241bp of *cat*-MuScr (scrambled sequence) in the intergenic region between *pstS* and *glmS* using Datsenko and Wanner method (2) |
| MuScr_origin 1_gyrA-FLAG | Mu Scrambled_origin1 *gyrA-FLAG-kan* | This work; P1 phage transduction of FLAG-*kan* at C-terminus of *gyrA* from Mu_origin 1_gyrA-FLAG |

| | | |
|---|---|---|
| MuScr_origin 2 | MG1655 *aptI-rsmG*::*MuScr* | This work; Integration of MuScr-*kan* in the intergenic region between *aptL* and *rsmG* using Datsenko and Wanner method and cured of *kan* using pCP20 (2) |
| MuScr_terminus 1 | MG1655 *yddY-yddW*::*MuScr* | This work; Integration of MuScr-*kan* in the intergenic region between *yddY* and *yddW* using Datsenko and Wanner method and cured of *kan* using pCP20 (2) |
| MuScr_terminus 2 | MG1655 *ydiY-pfkB*::*kan-MuScr* | This work; Integration of MuScr-*kan* in the intergenic region between *ydiY* and *pfkB* using Datsenko and Wanner method (2) |
| MG1655 Δ*recD* | MG1655 Δ*recD* | This work; P1 phage transduction of mutation from Keio collection followed by curing of the kanamycin resistance cassette (3) |
| PIR1 | F- Δ*lac169 rpoS(Am) robA1 creC510 hsdR514 endA recA1 uidA(*Δ*MluI)::pir-116* | Invitrogen; The *pir* gene in PIR1 encodes the replication protein $\pi$, which is required to replicate and maintain plasmids containing the R6Kγ origin. The strain is used to maintain pKD3_Mu, pKD4_Mu, pKD3_MuScr, pKD4_MuScr used in this study |
| MG1655 4Mu_gyrA-FLAG | Mu_ origin 1 *yddY-yddW*::*Mu ydiY-pfkB*:: *Mu aptI-rsmG*::*Mu gyrA-kan* | This work; Integration of Mu-*kan* using sequential P1 transduction of Mu_terminus 2, Mu_origin 2, and Mu_origin 1_gyrA-FLAG to Mu_terminus 1 strain with step-wise curing of the kanamycin resistance cassette(3) |
| **Plasmid** | **Description** | **Source** |
| pKD3 | Plasmid conferring chloramphenicol and ampicillin resistance | (2) |

| | | |
|---|---|---|
| pKD4 | Plasmid conferring kanamycin and ampicillin resistance | (2) |
| pKD3_Mu | Mu strong gyrase cleavage sequence cloned into pKD3 | This work |
| pKD3_MuScr | Mu Scrambled sequence cloned into pKD3 | This work |
| pKD4_Mu | Mu strong gyrase cleavage sequence cloned into pKD4 | This work |
| pKD4_MuScr | Mu Scrambled sequence cloned into pKD4 | This work |
| pKD4_gyrA-FLAG | C-terminus of FLAG-tagged *gyrA* cloned into pKD4 | This work |
| pCP20 | pCP20, repA101(ts), $Amp^R$ and $CAM^R$ | (2) |

**Supplementary Table S2. DNA oligonucleotides**

| Primers/gBlocks for plasmid construction | | |
|---|---|---|
| **Primer name** | **Sequence** | **Description** |
| Mu _Gibson_pKD3 | GATCTTCCGTCACAGGTAGGATGTGCTGC AAGGCGATTAAGTTGGGTAACGCCAGGG TTTTCCCAGTCACGACGTTGTAAAACGAC GGCCAGTGCCAAGCTTGCATGCCTGCAG GTCACTGGAGAAAGAAAGTGAAAGGAA GATAAAACGGGATTCATACACCGTTAAA TACCGGTTTAAAAATCCCGTGGCGCGTTT TAAAAAATCTGTGCGGGTGATTTTATGCC TGATTCTGTTTATTGCCTCAGAGCGGCGC TGACGCGTTTTCTGATGGCATCAAAAATT TCCTGTTCCCCGGTCTTATCCAGCCCCAT ATAAGGACGCGCAGGAACGCCTGCCGGG CGGGGTGCCATATCGGGTGTACCGCCCTC CCATGTCAGCCGTTAAGT | 389bp gBlock gene fragments containing the strong gyrase cleavage site Mu (4) for the construction of pKD3_Mu |
| MuScr_pKD3_1 | GATCTTCCGTCACAGGTAGGATGTGCTGC AAGGCGATTAAGTTGGGTAACGCCAGGG TTTTCCCAGTCACGACGTTGTAAAACGAC GGCCAGTGCCAAGCTTGCATGCCTGCAG GTCACTGGAGAAAGAAAGTGAAAGGAA GATAAAACGGGATTCATACACCGTTAAA TACCGGTTTAAAAATCCCGTGGCGCGTA GTCTTGATTTTAGAGTCTGAGCCGAGTGC GGTTTCATTTCCCGGGATAGTGGTTCAGA AACCTCTTTTTTCTGATGGCATCAAAAAT TTCCTGTTCCCCGGTCTTATCCAGCCCCA TATAAGGACGCGCAGGAACGCCTGCCGG | 389bp gBlock gene fragments containing the Mu scrambled sequence for the construction of pKD3_MuScr |

| | | |
|---|---|---|
| | GCGGGGTGCCATATCGGGTGTACCGCCC TCCCATGTCAGCCGTTAAGT | |
| pKD3_FWD | TCCCATGTCAGCCGTTAAGTG | Used in conjunction with pKD3_REV to amplify the backbone of pKD3 or pKD4_REV to amplify the backbone of pKD4 |
| pKD3_REV | CCTACCTGTGACGGAAGATCAC | Used in conjunction with pKD3_FWD to amplify the backbone of pKD3 |
| Cat_R | GCAACTGACTGAAATGCCTC | Used to confirm insertion of Mu or MuScr in pKD3 by Sanger sequencing |
| Cam_pKD3_ext_rev | AAGCAGAAGGCCATCCTGAC | Used to confirm insertion of Mu or MuScr in pKD3 by Sanger sequencing |
| cmR_int_fwd_1 | TCGTCTCAGCCAATCCCTG | Used to confirm insertion of Mu or MuScr in pKD3 by Sanger sequencing |
| Mu_ext_rev | CGAACTAAACCCTCATGGC | Used to confirm insertion of Mu or MuScr in pKD3 or pKD4 by Sanger sequencing |
| Mu_pKD4 | GAGGATATTCATATGGACCAATGTGCTG CAAGGCGATTAAGTTGGGTAACGCCAGG GTTTTCCCAGTCACGACGTTGTAAAACGA CGGCCAGTGCCAAGCTTGCATGCCTGCA GGTCACTGGAGAAAGAAAGTGAAAGGA AGATAAAACGGGATTCATACACCGTTAA ATACCGGTTTAAAAATCCCGTGGCGCGTT TTAAAAAATCTGTGCGGGTGATTTTATGC CTGATTCTGTTTATTGCCTCAGAGCGGCG CTGACGCGTTTTCTGATGGCATCAAAAAT TTCCTGTTCCCCGGTCTTATCCAGCCCCA TATAAGGACGCGCAGGAACGCCTGCCGG GCGGGGTGCCATATCGGGTGTACCGCCC TCCCATGTCAGCCGTTAAGT | 389bp gBlock gene fragments containing the strong gyrase cleavage site Mu (4) for the construction of pKD4_Mu |

| | | |
|---|---|---|
| MuScr_pKD4 | GAGGATATTCATATGGACCAATGTGCTG CAAGGCGATTAAGTTGGGTAACGCCAGG GTTTTCCCAGTCACGACGTTGTAAAACGA CGGCCAGTGCCAAGCTTGCATGCCTGCA GGTCACTGGAGAAAGAAAGTGAAAGGA AGATAAAACGGGATTCATACACCGTTAA ATACCGGTTTAAAAATCCCGTGGCGCGT AGTCTTGATTTTAGAGTCTGAGCCGAGTG CGGTTTCATTTCCCGGGATAGTGGTTCAG AAACCTCTTTTTTCTGATGGCATCAAAAA TTTCCTGTTCCCCGGTCTTATCCAGCCCC ATATAAGGACGCGCAGGAACGCCTGCCG GGCGGGGTGCCATATCGGGTGTACCGCC CTCCCATGTCAGCCGTTAAGT | 389bp gBlock gene fragments containing the Mu scrambled sequence for the construction of pKD4_MuScr |
| pKD4_SEQ_FWD | GCCATCACGAGATTTCGATT | Used to confirm insertion of Mu or MuScr by Sanger sequencing |
| gyrA_CFLAG | GAGGATATTCATATGGACCATTACTTGTC ATCGTCGTCCTTGTAGTCGGATCCTTCTT CTTCTGGCTCGTCGTCAACGTCCACTTCC GGAGCGATTTCATCGTCCCCTTCCGCTCC CATGTCAGCCGTTAAGTG | Gene fragments containing the FLAG tag to the C-terminus of GyrA; Used for insertion into pKD4 next to the *kan* cassette |
| pKD4_REV | TGGTCCATATGAATATCCTCCTTAG | Used in conjunction with pKD3_FWD to amplify pKD4 backbone |
| **Primers for chromosomal perturbations** | | |
| **Primer name** | **Sequence** | **Description** |
| cat_Mu_pstS_pkD3_FWD | TAAGCGTTGATATTCAGTCAATTACAAAC ATTAATAACGAGGAATAGGAACTTCATT TAAATGGCGCG | Used in conjunction with cat_Mu_glmS_pKD3_REV to amplify Mu or MuScr from pKD3_Mu or pKD3_MuScr for |

| | | |
|---|---|---|
| | | construction of Mu_origin 1 or MuScr_origin 1 |
| cat_Mu_glmS_pKD3_REV | CTTTTTCTCTGTCACAGAATGAAAATTTTTCTGTCATCTCTGGGCGGTACACCCGATATG | Used in conjunction with cat_Mu_pstS_pkD3_FWD to amplify Mu or MuScr from pKD3_Mu or pKD3_MuScr for construction of Mu_origin 1 or MuScr_origin 1 |
| gyrA_CFLAG_FWD | CAATTCAAACAAGGGAGATAGCTCCCTTTTGGCATGAAGAAGTAAAAGTGTAGGCTGGAGCTGCTTCG | Used in conjunction with gyrA_CFLAG_REV to amplify *gyrA-FLAG* from pKD4_gyrA-FLAG for construction of Mu_origin 1-gyrA-FLAG |
| gyrA_CFLAG_REV | GCGGAAGGGGACGATGAAATC | Used in conjunction with gyrA_CFLAG_FWD to amplify *gyrA-CFLAG* from pKD4_gyrA-FLAG for construction of Mu_origin 1-gyrA-FLAG |
| pKD4_rev_gyrA | GATGAAATCGCTCCGGAAGTGGACGTTGACGACGAGCCAGAAGAAGAATAATGGTCCATATGAATATCCTCCTTAG | Used in conjunction with gyrA_CFLAG_FWD to amplify *gyrA-kan* from pKD4 for construction of Mu_origin 1-gyrA-FLAGless |
| rsmG_Mu_FWD | GTTTTAATAAATGACATTTACACAACAAAAACCACCCATTGAGTGTAGGCTGGAGCTGCTTCG | Used in conjunction with rsmG_Mu_REV to amplify Mu or MuScr from pKD4_Mu or pKD4_MuScr for construction of Mu_origin 2 or MuScr_origin 2 |
| rsmG_Mu_REV | CTAAGAACCATCATTGGCTGTTAAAACATTATTAAAAATGGGGCGGTACACCCGATATG | Used in conjunction with rsmG_Mu_FWD to amplify Mu or MuScr from pKD4_Mu or pKD4_MuScr for construction of Mu_origin 2 or MuScr_origin 2 |
| yddW_Mu_FWD | ATATTCATACATTTTTATTAGGGATTTATGGCTGTTTAACGTGTAGGCTGGAGCTGCTTCG | Used in conjunction with yddW_Mu_REV to amplify Mu or MuScr from pKD4_Mu or pKD4_MuScr for construction of Mu_terminus 1 or MuScr_terminus 1 |

| | | |
|---|---|---|
| yddW_Mu_REV | CTAATAATCATGCTTACTTAAGTCAAATTAACCACACTTAGGGCGGTACACCCGATATG | Used in conjunction with yddW_Mu_FWD to amplify Mu or MuScr from pKD4_Mu or pKD4_MuScr for construction of Mu_terminus 1 or MuScr_terminus1 |
| pfkB_Mu_FWD | GTATTCTTATTTCATTTTTTGAATAAGCATGTGGCGAAAACAGTGTAGGCTGGAGCTGCTTCG | Used in conjunction with pfkB_Mu_REV to amplify Mu or MuScr from pKD4_Mu or pKD4_MuScr for construction of Mu_terminus 2 or MuScr_terminus 2 |
| pfkB_Mu_REV | GAGCTTTATTTAAAATTTTGCAGATAAATATATATAAATAAAAATCGGGCGGTACACCCGATATG | Used in conjunction with pfkB_Mu_FWD to amplify Mu or MuScr from pKD4_Mu or pKD4_MuScr for construction of Mu_terminus 2 or MuScr_terminus 2 |
| **Primers used to verify gene perturbations** | | |
| **Primer name** | **Sequence** | **Description** |
| pstS-int-fwd | AGGCTTGCTTCTGCAAACAC | Used in conjunction with glmS-int-rev to confirm chromosomal location of Mu or MuScr in Mu_origin 1and MuScr_origin 1, respectively |
| glmS-int-rev | GGTGATTGCACCGATCTTCT | Used in conjunction with pstS-int-fwd to confirm chromosomal location of Mu or MuScr in Mu_origin 1 and MuScr_origin 1, respectively |
| cat_R | GCAACTGACTGAAATGCCTC | Used in conjunction with Mu_int_rev to confirm chromosomal integration of Mu or MuScr |
| Mu_int_rev | CGCGCCACGGGATTTTTAAACC | Used in conjunction with cat_R to confirm chromosomal integration of Mu or MuScr |

| | | |
|---|---|---|
| rsmG_ext_fwd | CAAACAATAAGTAGCCAAAAG | Used in conjunction with rsmG_int_rev to confirm chromosomal location of Mu or MuScr in Mu_origin 2 and MuScr_origin 2, respectively |
| rsmG_int_rev | AGCCCATTTCACTCTGTTGG | Used in conjunction with rsmG_ext_fwd to confirm chromosomal location of Mu or MuScr in Mu_origin 2 and MuScr_origin 2, respectively |
| yddW_ext_fwd | ATACTGAAAAGAAATAAGCG | Used in conjunction with yddW_int_rev to confirm chromosomal location of Mu or MuScr in Mu_terminus 1and MuScr_terminus 1, respectively |
| yddW_int_rev | TAAAAGCACGCCTCCAGAGT | Used in conjunction with yddW_ext_fwd to confirm chromosomal location of Mu or MuScr in Mu_terminus 1and MuScr_terminus 1, respectively |
| pfkB_ext_fwd | TGGTGTCAGCCGTAAGTGAG | Used in conjunction with pfkB_int_rev to confirm chromosomal location of Mu or MuScr in Mu_terminus 2 and MuScr terminus 2, respectively |
| pfkB_int_rev | ACCAGCGCACTGAGTTCTTT | Used in conjunction with pfkB_ext_fwd to confirm chromosomal location of Mu or MuScr in Mu_terminus 2 and MuScr_terminus 2, respectively |
| recD_int_fwd | TGCGCTTCTGTTGCATAAAC | Used in conjunction with recD_int_rev to confirm deletion of *recD* |
| recD_int_rev | TTTACAGAGCGGCGAAGATT | Used in conjunction with recD_int_fwd to confirm deletion of *recD* |
| argA_int_fwd | TCAAGGGGTGAAGTTCTGCT | Used in conjunction with recB_int_rev to confirm deletion of *recD* and the excision of *kan* cassette |

| | | |
|---|---|---|
| recB_int_rev | TGAGATGTTTGCCGGTATGA | Used in conjunction with argA_int_fwd to confirm deletion of *recD* and the excision of *kan* cassette |
| kan_int_rev | ATGATGGATACTTTCTCGGCAGGAG | Used to confirm insertion of *FLAG-kan* or *kan* to the C-terminus of *gyrA* by Sanger sequencing |
| gyrA_ext_seq_rev | CCAAACTTTACCGTGCCCTA | Used in conjunction with gyrA_int_fwd4 to confirm insertion of *FLAG-kan* or *kan* to the C-terminus of *gyrA* |
| gyrA_int_fwd4 | GGCGATAAAGTCGTCTCTCTGA | Used in conjunction with gyrA_ext_seq_rev to confirm insertion of *FLAG-kan* or *kan* to the C-terminus of *gyrA* |
| **Primers for ChIP-qPCR analysis** | | |
| **Site** | **Forward primer (5′→3′)** | **Reverse Primer (5′→3′)** |
| Mu | CGACGTTGTAAAACGACGGCC | CGCGCCACGGGATTTTTAAACC |
| *trkH* | CTTTACCAGTATGAACCCGGTGG | CCAGAAAGTCGGGGTAAAGAGC |
| *mobA* | CTCACCAGCTTCCTGCTGC | ATTGAAGATTCACTGGCGGATTACC |
| *nuoN* | AGAGATCAGCACAACGATAC | CTACCTGCGCGTGGCGGTGA |

**Supplementary Table S3. Macrodomain boundary**

| **Macrodomain**(5, 6) | **Start*** | **End*** |
|---|---|---|
| NS-right | 46418 | 603415 |
| Right | 603416 | 1206830 |

| | | |
|---|---|---|
| Ter | 1206831 | 2181576 |
| Left | 2181577 | 2877824 |
| NS-left | 2877825 | 3759738 |
| Ori | 3759739 | 46417 |

* MD coordinates were adjusted to Mu-origin 1 genome
